# Programmable DNA shell scaffolds for directional membrane budding

**DOI:** 10.1101/2024.01.18.576181

**Authors:** Michael T. Pinner, Hendrik Dietz

## Abstract

In the pursuit of replicating biological processes at the nanoscale, controlling cellular membrane dynamics has emerged as a key area of interest. Here, we report a system mimicking virus assembly to control directional membrane budding. We employ three-dimensional DNA origami techniques to construct cholesterol-modified triangles that self-assemble into polyhedral shells on lipid vesicles, resulting in gradual curvature induction, bud formation and spontaneous neck scission. Strategic positioning of cholesterols on the triangle surface provides control over the directionality of bud growth and yields daughter vesicles with DNA endo- or exoskeletons reminiscent of clathrin-coated vesicles. This process occurs with rapid kinetics and across various lipid compositions. When combined into a two-step process, nested bivesicular objects with DNA shells encapsulated between lipid vesicles could be produced. Our work replicates key aspects of natural endocytic and exocytic pathways, opening new avenues for exploring membrane mechanics and applications in targeted drug delivery and synthetic biology.

## Main text

Membrane budding is a ubiquitous biological phenomenon in which parent lipid membranes release daughter vesicles for the transport of materials across the membrane. In biological systems, membrane curvature changes are often induced by curved proteins such as clathrin. With its distinctive triskelion shape, clathrin acts as a molecular scaffold by binding to membrane-bound receptor-adaptor protein complexes, followed by self-assembly into spherical cage-like structures that induce curvature in the cell membrane^1–3^. Recent studies confirmed clathrin’s membrane-bending properties in vitro^4^ and in cells^5^. We hypothesised that any material capable of self-assembly into curved shapes could act as a molecular scaffold for membrane budding, with DNA origami emerging as a promising candidate. Herein, a several kilobases long, single-stranded ‘scaffold’ strand is mixed with an array of shorter oligonucleotides and exposed to a thermal annealing ramp to drive the folding of the DNA origami structure^6–8^. Previous efforts in the field related to lipid membranes have successfully tethered membrane-spanning DNA nanopores of various complexity to lipid bilayers using hydrophobic moieties such as cholesterol^9^, ethyl-phosphorothioate^10^ or tetraphenylporphyrin^11^. Notably, cholesterol-labelled DNA origami objects tethered to lipid membranes retain diffusive mobility influenced by factors like linker oligonucleotide length^12^, paving the way for the 2D assembly of monomeric structures into multimeric lattices^13^. Prior studies have also shown that both curved^14–16^ and planar^14^ origami structures can induce tubulation in giant unilamellar lipid vesicles. On-membrane polymerisation of origami structures likewise resulted in the deformation of the underlying membrane reminiscent of clathrin networks involved in cellular budding processes^13,17–20^.

Building upon this foundational work, we have developed an artificial, DNA-origami-based membrane budding system with spontaneous neck scission. At its core are three-dimensional membrane-interacting triangular subunits constructed from DNA with bevelled edges^21^. Strategically placed shape-complementary protrusions and recesses at the triangle edges direct the self-assembly of these triangular subunits into icosahedral shells through adhesive base pair stacking interactions^22^. Interactions with lipid membranes of giant vesicles (GVs) were promoted using cholesterol moieties placed on the origami structure. We hypothesised that membrane deformation processes may be sufficient for the spontaneous scission of the bud neck without the need for active constriction by GTPases like dynamin^23^. Indeed, the self-assembly of membrane-bound triangles induced local curvature and finally led to the formation and eventual scission of buds. The positioning of cholesterol determined whether the bud-formation occurs inward or outward from the source vesicle, leading to DNA-shell-coated vesicles (DCV) and vesicle-coated DNA shells (VCD), respectively.

### DNA origami triangles as molecular scaffolds

We adapted a previously developed programmable icosahedral shell canvas, in which twenty equilateral DNA origami triangles form a closed icosahedron triggered by elevated concentrations of magnesium chloride for membrane-supported assembly (Supplementary Figure 1 & Supplementary Tables 1-24)^21^. To provide the necessary hydrophobic interactions required for association with lipid membranes, we incorporated multiple cholesterol-bearing oligonucleotides (chol-oligos) at the shell-inward-facing surface of the constituent triangular DNA origami subunits (Fig. 1a & Supplementary Figure 2). We hypothesised that the 2D diffusion of triangles on a fluid membrane can support the assembly of icosahedral shells whilst pulling membrane material along, resulting in increasingly deformed membrane buds growing away from the parent membrane until the bud neck is cut and a DNA-shell-coated vesicle (DCV) is released into solution.

**Fig. 1.**
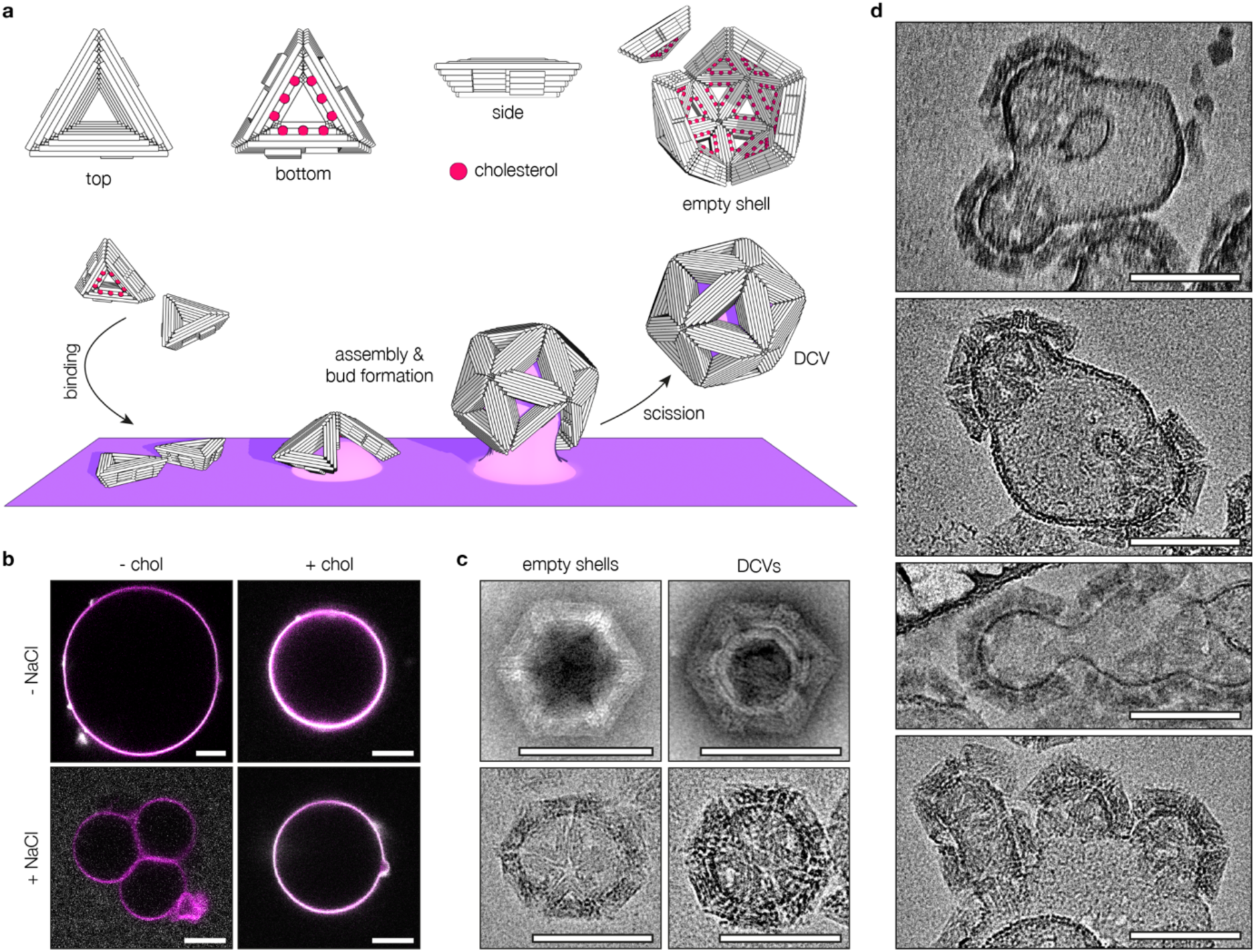
DNA-shell-coated vesicles form by membrane budding. **a,** Schematic of a DNA origami triangle-based membrane budding system. The triangular structures are decorated with cholesterol moieties to tether them to the membrane of a lipid vesicle. Upon increasing the MgCl_2_ concentration, the triangles self-assemble into polyhedral shells, pulling the membrane underneath along to form a bud. Following spontaneous neck scission, a DNA-shell-coated vesicle (DCV) is released. **b,** Pseudocoloured micrographs of DNA origami triangle attachment (grey) onto giant vesicles (GVs, magenta). Triangles will adsorb onto GVs even if they are not decorated with cholesterol (top left & top right panels). However, adding sodium chloride suppresses this non-specific adsorption (bottom left panel), and membrane binding will only occur if cholesterol is attached to the triangle (bottom right panel). Scale bars: 10 µm. **c,** Transmission electron micrographs of empty DNA shells and DCVs. In negative stain TEM (top panels), lipid vesicles within DCVs appear as white rings inside the DNA shell. In cryogenic electron microscopy (cryoEM, bottom panels), the vesicle is visible as a dark ring inside the DNA shell. Scale bars: 100 nm. **d,** CryoEM micrographs of large vesicle (LV) deformation upon assembly of membrane-bound DNA origami triangles. Bud curvature increases gradually and closely follows the intrinsic curvature of the assembling shells. Scale bars: 100 nm.

We screened cholesterol density and positioning to balance the solubility of the triangles with membrane affinity (Supplementary Figure 3a & b) and then validated their assembly into the expected icosahedral shells in the absence of membranes (Supplementary Figure 3c). Introducing cholesterol resulted in a shift from the previously observed relatively uniform icosahedral shells to a mixture containing both octahedral and icosahedral shells. This variation was controllable: by limiting the reach of each cholesterol moiety, we could largely revert the assembly to predominantly icosahedral structures. The addition of lipid membranes into the system introduced another variable, again leading to the occurrence of octahedral structures. This tendency varied depending on the amount of cholesterol per triangle and the presence of lipid vesicles, detailed in Supplementary Figures 4, 5 & 6a. We believe this effect arises from a complex interplay of forces, including crowding of membrane-bound triangles, altered or biased relative alignment of triangles (possibly influenced by local curvature induction), and elastic deformation within the shells caused by less-than-ideal angles between the triangle subunits. Our shells share this structural ambiguity with clathrin, which likewise assembles into differently shaped cages without negatively impacting its role as a molecular scaffold, and we thus considered both shell types suitable for our objectives^3^.

We prepared giant vesicles (GVs) composed entirely of DOPC and fluorescently labelled DOPE to serve as model membranes in our budding assays^24^. Aliquots of GVs and chol-decorated triangles were then mixed to anchor the nanostructures to the vesicles (Fig. 1b, detailed in Supplementary Figure 7). The inclusion of NaCl in the reaction mixture minimised non-specific binding and ensured that the triangles attached to the vesicles in the correct orientation as defined by the positioning of their cholesterol moieties^16^. We then triggered the assembly of membrane-bound triangles by increasing the MgCl_2_ concentration and incubating the solution at 37 °C for up to multiple days. Negative stain transmission electron microscopy (TEM) imaging revealed abundant DCVs, characterised by dark DNA shells fully encapsulating bright lipid vesicles (Fig. 1c, top right; Supplementary Figure 8). Occasionally, the shell-enclosed vesicles appeared deformed or cracked, which we attribute to staining and drying artefacts (Supplementary Figure 9). To also study DCVs in their native state, we acquired micrographs of vitrified samples by cryogenic transmission electron microscopy (cryoEM, Fig. 1c, bottom right; Supplementary Figure 10a-c). This data generally agrees with the conclusions drawn from negative stain TEM data, but the enclosed vesicles seen by cryoEM rarely had any defects except for occasionally encapsulating even smaller vesicles themselves. In the cryo-EM micrographs, the vesicle typically fills up the available space inside the origami shell, with the membrane in direct contact with the enclosed DNA shell.

As the size difference between DCVs and GVs spans multiple orders of magnitude, direct observation of DCV formation is challenging. To overcome this problem, we instead used large vesicles (LVs) of approx. 200 nm in our budding assays. By additionally using isotonic MgCl_2_ solution and shortening the time of shell assembly before vitrification, we could resolve intermediate states of DCV formation by cryoEM (Fig. 1d & Supplementary Figure 10d). Whilst membrane patches free of DNA triangle subunits appear undisturbed, those with triangles on them had deformations that matched the curvature of the triangle assemblies. Notably, some vesicles carry multiple buds with clearly visible necks. This image data strongly supports the notion that the assembly of membrane-bound triangles induces curvature and that budding is the mechanism behind DCV formation.

### Characterisation of DCVs

We analysed DCV formation efficiency by agarose gel electrophoresis using fluorescently labelled DNA origami and GVs (Supplementary Figure 11). Lipid material colocalised exclusively with the higher-order shell assemblies and was virtually absent at the level of monomeric triangles and intermediates. We also note that cholesterol-free triangles, variants incapable of shell formation, and triangles added to GVs after several hours of membrane-free assembly all failed to produce DCVs efficiently (Supplementary Figures 6b, 12 & 13). These findings underscore the critical role of on-membrane triangle assembly in DCV formation.

To evaluate whether the lipid vesicles were fully enclosed by well-ordered polyhedral DNA shells, we recorded single-axis tilt series to generate tomograms, allowing us to study DCVs in 3D (Fig. 2a). The representative tomograms confirm full engulfment of the vesicle by closed shells featuring the symmetry properties expected by design. Occasionally, we found DCVs where the DNA origami shell had assembly “scars” consisting of one to several missing triangles (Fig. 2b). Considering that the increasingly high curvature at the bud neck can become a steric barrier to completion of the DNA origami shell, we believe the occurrence of scars to be a signature of the proposed budding process. In this scenario, spontaneous membrane scission outcompetes the completion of the DNA origami shell.

**Fig. 2.**
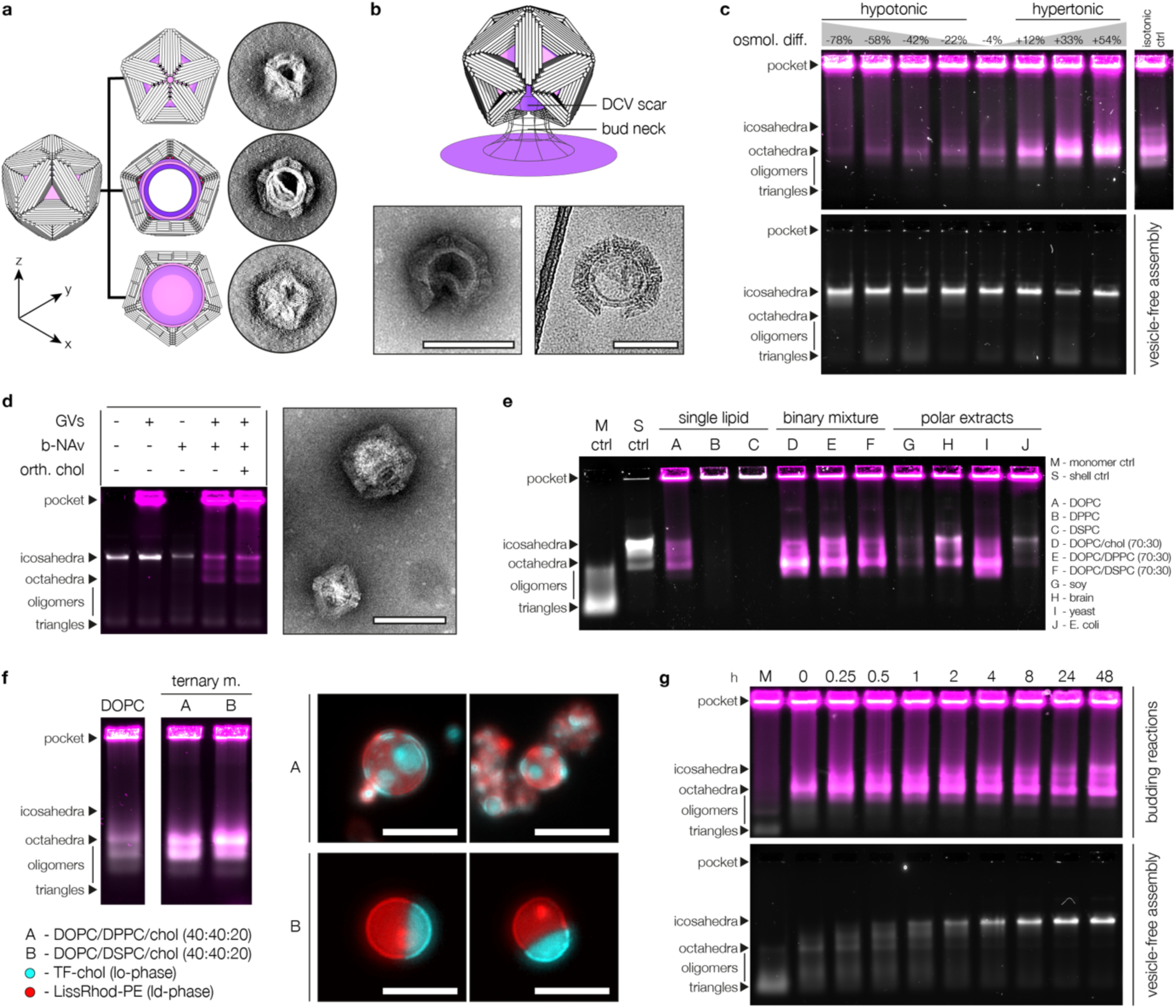
Characterisation of DNA-shell-coated vesicles. **a,** Tomogram of an icosahedral DCV depicted as summed up z-slices of the top, central and bottom segments. The topmost segment appears as a tilted, incomplete pentamer. The central segment reveals a lipid vesicle trapped within the origami shell. The bottommost segment is a pentameric shell cap. **b,** DCVs occasionally appear incomplete with visible ‘scars’ of missing triangles. The bud neck prevents closure of the DNA shell by steric hindrance, resulting in a gap in the origami shell that remains visible after scission of the neck. Left panel: negative stain TEM; Right panel: cryoEM. Scale bars: 100 nm. **c,** Agarose gel of DCVs prepared under varying osmotic conditions. Samples in the top gel contain triangles (grey, EtBr) and GVs (magenta, DOPE-Atto643), whereas the bottom gel contains triangles only and serves as assembly control. Glycine and MgCl_2_ were added at increasing concentrations to vary sample tonicity and promote triangle assembly. Budding efficiency strongly correlates with tonicity, working best under hypertonic conditions and coming to a near halt at the most hypotonic condition tested. The unhindered assembly of triangles in absence of vesicles indicates that DCV formation is strongly influenced by changes in membrane tension. **d,** Left: Agarose gel of DCVs prepared by using biotin-NeutrAvidin (b-NAv) interactions to connect origami triangles (grey, Atto643) and GVs (magenta, DOPE-Atto488). The addition of orthogonal chol-oligos did not influence DCV yields. Right: TEM micrographs of DCVs obtained by this approach. Scale bar: 100 nm. **e,** Agarose gel of DCVs obtained from GVs of varying lipid composition. Only GVs composed entirely of high-melting lipids prevented DCV formation. Grey: DNA (EtBr); Magenta: Lipids (DOPE-Atto643); **f,** Left: Agarose gel comparing DCVs obtained from GVs composed of DOPC or phase-separated ternary lipid mixtures. Phase-separated vesicles yielded more DCVs, suggesting an influence of phase boundaries on the budding mechanism. Grey: DNA (EtBr); Magenta: Lipids (LissRhod-PE); Right: Micrographs of exemplary phase-separated vesicles showing the lipid-ordered (cyan) and -disordered (red) phases. Scale bars: 20 µm. **g,** DCV formation kinetics (top) compared against vesicle-free assembly kinetics. DCV budding occurs instantly and yields mostly octahedral species, but at longer incubation times a notable fraction of icosahedral DCVs is formed. Grey: DNA (EtBr); Magenta: Lipids (DOPE-Atto643);

The addition of concentrated MgCl_2_ solution as a trigger for shell assembly inevitably increases the osmolality of the sample buffer and may thus alter the biophysical properties of the membrane by deflating the vesicles. Indeed, hyperosmotic stress has previously been exploited to alter vesicle shape by membrane-bound DNA origami^16^. We studied the influence of membrane tension on DCV yield by mixing triangle-covered GVs with increasing amounts of glycine as an osmolyte. Interestingly, DCV yields correlated strongly with the osmotic environment. In the hypotonic regime, we observed moderate to low yields, with the lowest tonicity tested producing only a very faint band by agarose gel electrophoresis (Fig. 2c, Supplementary Figure 14 & Supplementary Table 25). Conversely, as we transitioned into the hypertonic regime, we noted a marked and progressive increase in DCV yields, with the most pronounced yield achieved at the highest buffer osmolality. These findings reveal a direct relationship between membrane tension and DCV yields, reinforcing the notion that DCVs emerge from intricate membrane remodelling processes.

Our system’s sensitivity to membrane tension draws a parallel to clathrin-mediated endocytosis, which depends on actin engagement for budding under increased membrane tension^25^. Another point of similarity is that the assembly of both our DNA triangles and clathrin itself is sufficient to deform model membranes in vitro^4^. However, vesicle deformation has also been demonstrated by attaching DNA origami structures to lipid vesicles using cholesterol as an anchor, not requiring the assembly of subunits^14^. Having demonstrated the necessity of triangle assembly for DCV formation (Supplementary Figure 12), we next investigated the roles of cholesterol insertion and outer leaflet expansion in inducing curvature and budding. To evaluate this, we prepared GVs using a lipid mixture containing 3% biotinylated DOPE and replaced chol-oligos with biotinylated oligonucleotides linked to NeutrAvidin (b-NAv) as membrane anchors. This allowed DNA triangles to attach to the membrane via NeutrAvidin-biotin interactions without inserting foreign lipids into the outer leaflet. Remarkably, this approach still resulted in DCV formation and adding excess chol-oligos with an orthogonal sequence did not improve yields (Fig. 2d & Supplementary Figure 15). These findings show that cholesterol insertion is not required for DCV formation, supporting the idea that membrane buds are primarily shaped by shell assembly.

In our study, we primarily used vesicles composed of DOPC, supplemented with a small fraction of fluorescently labelled DOPE species. The high fluidity and homogeneity of DOPC membranes provide idealised conditions that do not necessarily reflect the properties of biological membranes. We next investigated the versatility of our budding system using GVs with diverse lipid compositions. These included single-lipid species compositions (low-melting DOPC, and high-melting DPPC and DSPC), binary mixtures (70% DOPC with either 30% cholesterol, DPPC, or DSPC), and four polar extracts of natural origin (derived from soy, brain, yeast, and *E. coli*). With the exception of GVs composed entirely of high-melting lipids, DCVs successfully formed from all tested compositions with varying efficiencies (Fig. 2e & Supplementary Figure 16). The successful formation of DCVs from binary mixtures containing high-melting lipids suggests that the overall membrane fluidity, rather than the mere presence of high-melting lipids, is a critical factor in determining budding success. This is supported by the fact that low-melting DOPC constituted the majority (70%) of these mixtures, rendering the bilayer overall fluid.

Another important factor for DCV formation is neck scission. Constriction of the bud neck may be driven by spontaneous and induced curvature at the neck due to a high triangle density on the membrane and adhesion forces^26,27^. Scission can also be helped by line tension in phase-separated membranes, where the phase boundary represents a linear defect along which fission is facilitated^28^. Indeed, when comparing DCV yields obtained from pure DOPC membranes versus phase-separated membranes composed of DOPC/DPPC/chol or DOPC/DSPC/chol (40:40:20 mol%), we found that the phase-separated membranes generally yielded more DCVs (Fig. 2f & Supplementary Figure 17).

We next analysed the kinetics of DCV formation by starting the assembly reaction and collecting aliquots at various time points. The budding process was rapid, with a substantial number of DCVs already present at the 0-minute time point (Fig. 2g & Supplementary Figure 18). We attribute the rapid kinetics to the membrane confinement of the origami triangles, which generates elevated local concentrations and restricts mobility to two dimensions. Consistent with earlier observations, the majority of DCVs formed initially exhibited an octahedral geometry. Over time, however, a growing fraction of icosahedral DCVs was detected. This progression mirrors the behaviour observed in vesicle-free control assemblies, where DNA origami triangles rapidly assemble into intermediates resembling octahedra in the agarose gel, which subsequently mature into icosahedral shells. However, the maturation of octahedral DCVs into icosahedral ones appears restricted. We believe that size constraints imposed by the vesicle inside the DCV stabilises its configuration and limits the expansion of the inner cavity necessary to form larger icosahedral species.

### Inward budding

Having studied the formation and characterisation of DCVs, we next investigated the directional reversal of this process. DCV budding mimics endocytic processes with the important difference that in our system the triangular subunits approach the parent vesicle from the exterior. By shifting the placement of cholesterols on the triangle surface from the bottom (shell-inward) to the top (shell-outward) face, we hypothesised that we could reverse the budding directionality in a process reminiscent of exocytosis. As the triangles would still bind to the vesicles from the outside, we expect the formation and release of vesicle-coated DNA shells (VCDs) into the lumen of the parent vesicle (Fig. 3a), effectively creating an endosome-like compartment with a DNA origami endoskeleton. We prepared large vesicles (LVs) by extruding GVs through 200 nm-sized pore filters to facilitate cryoEM imaging, mixed them with triangles carrying one cholesterol moiety per side and triggered shell assembly. Direct imaging with cryoEM (Fig. 3b) revealed VCDs as lipid vesicles tightly wrapped around icosahedral shells located within their respective parent vesicles. Occasionally, we also observed free VCDs, presumably set free by the bursting of the parent vesicle, and the presence of holes in the DNA shell similar to those observed in DCVs.

**Fig. 3.**
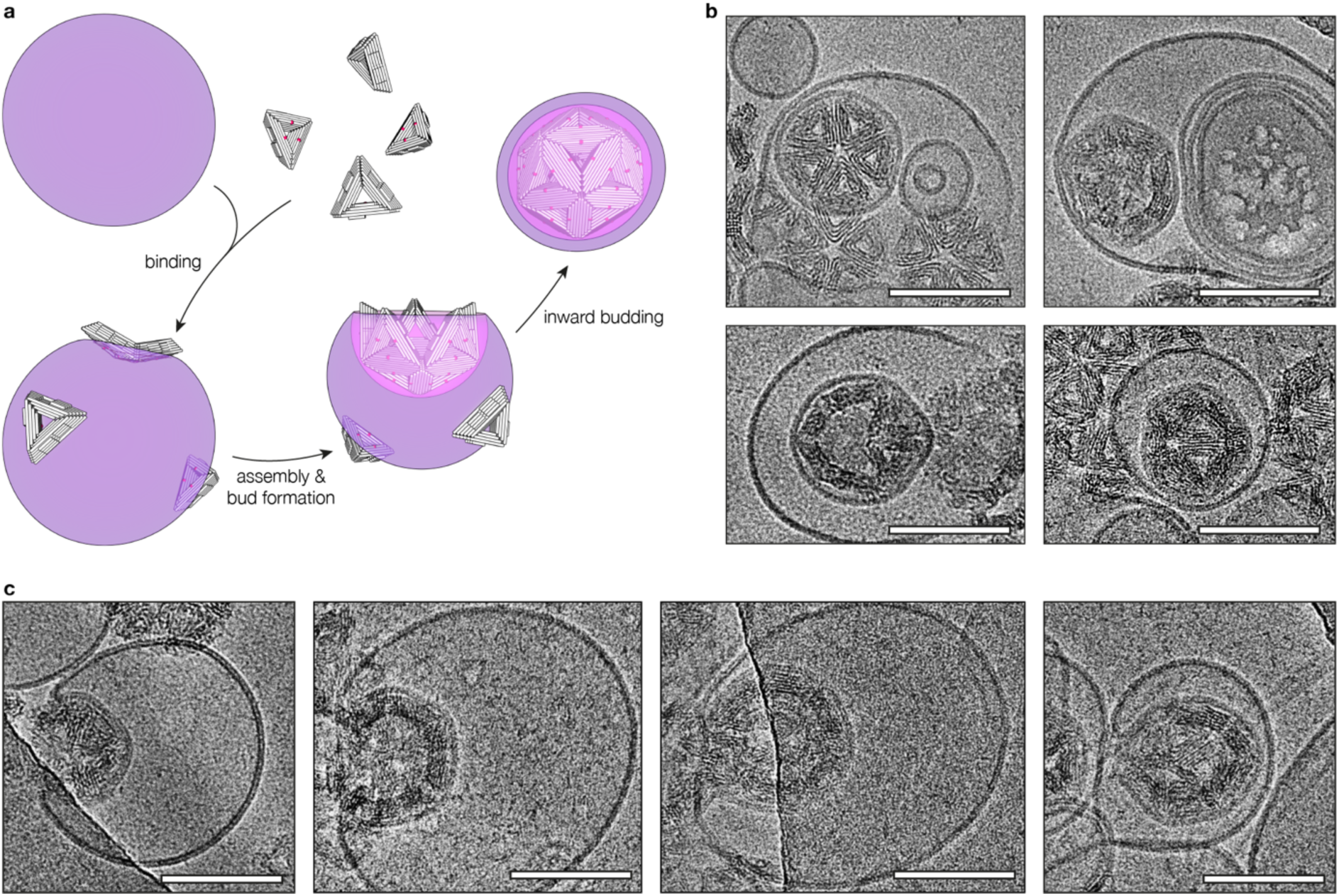
Vesicle-coated DNA shells. **a,** Illustration of vesicle-coated DNA shell (VCD) formation by budding into its parent vesicle. The budding direction is reversed with respect to DCVs by shifting the cholesterol linker handles from the shell-inner face to the shell-outer face of the triangle subunits. **b,** cryoEM micrographs of VCDs within their parent vesicles. Scale bars: 100 nm. **c,** cryoEM micrographs of inward-oriented membrane buds at different stages. Scale bars: 100 nm.

To capture the dynamics of the suspected budding process, we prepared vitrified samples with shortened assembly times and analysed them by cryoEM. The micrographs we obtained revealed the gradual deformation of the parent vesicle in response to the assembly of membrane-bound triangles, leading to the formation of inward-growing buds (Fig. 3c). In contrast to fully formed VCDs, these particles remain connected to the parent membrane, sometimes just by a narrow bud neck.

We finally explored the creation of bivesicular shell structures by combining outward and inward budding reactions in a two-step process using triangles with sequence-orthogonal linker handles on both faces (Fig. 4a). First, we produced DCVs by hybridising chol-oligos to the triangles’ bottom face and mixing the triangles with GVs as described earlier. After isolating the DCVs by centrifugation, we prepared LVs and coated them with chol-oligos complementary to the linkers on the DCV’s outer face. Mixing the DCV concentrate with these LVs and incubating at 37 °C resulted in the formation of vesicle-coated DCVs (VCDCVs), as confirmed by cryoEM (Fig. 4b). Like VCDs, VCDCVs were typically enclosed within their parent vesicles as a result of the proposed budding mechanism. These particles may be considered simple examples of synthetically created organelles featuring a cytoplasm-like compartment (the inner vesicle) and a periplasm-like compartment (the void between the outer envelope membrane and the inner lipid membrane). The icosahedral DNA shell acts as a stabilising mechanical skeleton, offering engineering options for including additional molecular functionalities.

**Fig. 4.**
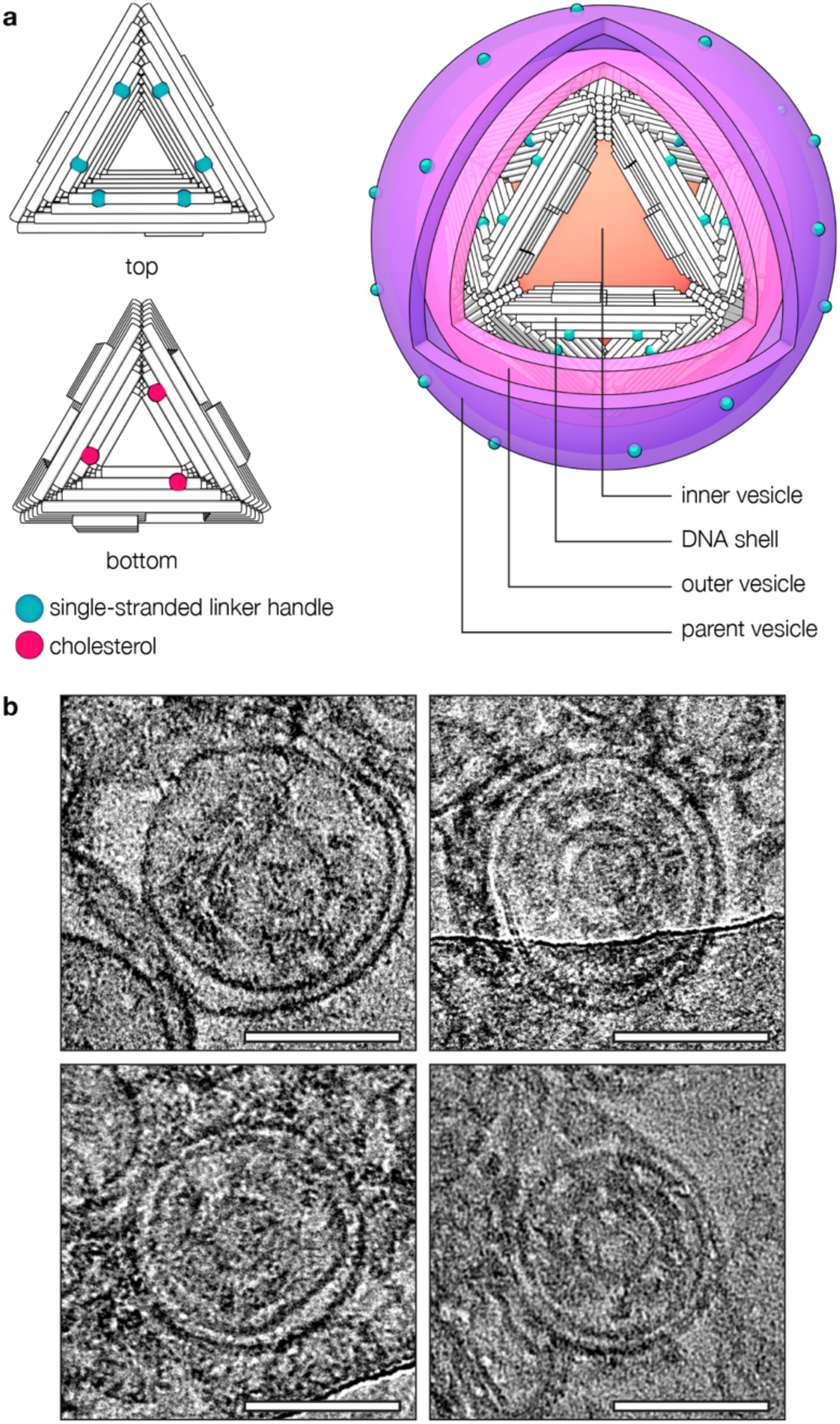
Multivesicular DNA shells. **a,** Illustration of a bivesicular DNA shell (vesicle-coated DCV; VCDCV) contained within its parent vesicle, produced in a 2-step budding assay using triangles with orthogonal linker handles on their top and bottom faces. In the first step, the bottom face linkers are hybridised to cholesterol-modified DNA strands for DCV formation by outward budding. Then, DCVs are mixed with LVs carrying single-stranded linker handles complementary to those on the shell-outer face of the triangles, resulting in inward budding and VCDCV formation. As a result, VCDCVs have four layers: An inner vesicle stemming from the first step (orange), the DNA origami shell (white), an outer vesicle (pink) coupled to the shell by complementary linker handles, and the parent vesicle (purple) with its DNA linker handles (green blobs). **b,** cryoEM micrographs of VCDCVs. Images were Gaussian blurred and levelled for clarity. Scale bars: 100 nm.

## Discussion

In this study, we developed a DNA origami-based membrane budding system that recapitulates key aspects of clathrin-mediated endocytosis (CME) without relying on components of cellular budding machineries. Like clathrin, the DNA origami triangles function as molecular scaffolds whose self-assembly into cage-like structures induces membrane deformation and ultimately drives the formation of vesicles. Free energy is supplied by the self-assembly process and relayed onto the bilayer by either cholesterol or biotin-NeutrAvidin linkages, mimicking the way clathrin associates with membranes via adaptor proteins. Using average stacking energies reported for DNA nanostructures (−2.97 k_B_T, T = 300 K), we estimate the free energy of icosahedral assembly (30 edges with 16 base-stacking interactions each) at approx. −1400 k_B_T, and −570 k_B_T for octahedra^29^. Our estimates exceed the energetic cost of bending DOPC membranes into spherical vesicles (460 k_B_T), derived from Helfrich’s theory (E = 8πk_C_, where k_C_ = 18.3 k_B_T at 298 K), and support our experimental observations^30,31^.

Both clathrin cages and DNA origami shells demonstrate structural flexibility, forming various geometries while maintaining their scaffolding function^3^. The triangles noticeably shifted from icosahedral to octahedral shells when assembled on the lipid membrane with influencing factors including lipid composition and membrane tether type. We suspect electrostatic interactions between the membrane and the origami, as well as molecular crowding effects, as key drivers behind this phenomenon. However, a detailed analysis lies beyond the scope of this study.

Our system demonstrates robust performance across a variety of lipid compositions, including natural lipid extracts, with rapid kinetics highlighting its versatility and potential for broader applications. Nonetheless, we identified limitations under specific conditions. DCV formation was impaired in membranes composed entirely of high-melting lipids (DPPC, DSPC) and yields were reduced under hypotonic conditions. These constraints can be explained by the Helfrich Hamiltonian, which relates the energetic cost of membrane deformation to the mechanical properties of the bilayer^30^. Higher bending rigidities of gel-phase membranes and increased membrane tension under hypotonic conditions impose energetic penalties directly impacting DCV yields^32^. While clathrin underlies the same limitations, biological membranes make use of the cytoskeleton to enable CME even under high membrane tension^25^. Future work could explore active neck constriction to overcome the current limitations of our system.

In summary, we present a versatile platform for engineering vesicles of controlled size featuring addressable endo- or exoskeletons. This approach can be extended to encapsulate molecules of interest during the budding process, potentially leading to molecular payload applications. Using nested bivesicular structures, it becomes possible to design scenarios where two different molecular payloads are delivered simultaneously but separately. This work may thus pave the way for innovative applications in targeted drug delivery, vaccine development, and synthetic biology. The potential to manipulate membrane dynamics at the nanoscale also opens new horizons in synthesising artificial cells and organelles, offering a valuable tool for researchers in their quest to harness the power of nanotechnology for practical and therapeutic purposes.

## Methods

### Materials

All chemicals were purchased from Sigma Aldrich and Carl Roth unless stated otherwise. 1,2-dioleoyl-sn-glycero-3-phosphocholine (DOPC), 1,2-dipalmitoyl-sn-glycero-3-phosphocholine (DPPC), 1,2-distearoyl-sn-glycero-3-phosphocholine (DSPC), polar lipid extracts (soy, brain, yeast, e. coli), TopFluor cholesterol and Lissamine Rhodamine B-1,2-dioleoyl-sn-glycero-3-phosphoethanolamine (DOPE) were obtained from Avanti Research. Atto488- and Atto643-modified DOPE was obtained from Atto-Tec. NeutrAvidin was purchased from Thermo Scientific. Regular DNA oligonucleotides were purchased from IDT, and modified oligonucleotides were obtained from Biomers. Scaffold DNA was produced biotechnologically in-house. Buffer osmolalities were measured using an Osmomat 3000 freeze point osmometer (Gonotec). For buffer compositions, refer to Supplementary Table 26.

### DNA origami folding, purification & quantification

DNA origami triangles were folded as described previously^21^. Briefly, circular scaffold strands of bacteriophage origin (sc8064, 50 nM; Supplementary Table 1) and staple oligonucleotides (200 nM per staple; Supplementary Tables 2-23) were mixed in folding buffer and exposed to a thermal annealing ramp (65 °C, 15’; 58-53 °C, 1 h/°C; stored at 20 °C) in a Tetrad2 (Bio-Rad) thermocycler. For some experiments, structures were fluorescently labelled by including Atto643 labelled oligonucleotides in the folding reaction. The folded product was purified by agarose gel electrophoresis by excising the leading band, crushing the gel pieces and centrifuging them in 0.45 µm Costar Spin-X spin columns (Corning) for 10 min at 8 krcf. When needed, the purified structures were washed and concentrated by ultrafiltration (Amicon Ultra 0.5 mL, 100 kDa cut-off, Millipore), exchanging the buffer for sodium buffer. Origami solutions were quantified using a NanoDrop8000 spectrophotometer (Thermo Scientific).

### Vesicle preparation and quantification

Giant vesicles were prepared by gentle swelling of dry lipid films on polyvinyl alcohol gels as described previously^24^, with slight modifications. A petri dish (⌀ 10 cm) was plasma cleaned in a glow discharge device, and then 1800 µl of a 5% solution of polyvinyl alcohol (Mowiol 28-99, Sigma Aldrich) in ddH_2_O was evenly distributed across its surface. The dish was incubated for at least 30 min at 50 °C until a dry gel formed. Then, 162 µl of the lipid mixture (typically DOPC with additional 0.5-1% fluorescently labelled DOPE if visualisation by fluorescence was desired, or 0.4% TopFluor cholesterol and LissRhod-PE each for visualisation of lipid phases. Total lipid concentration: 2.54 mM) in chloroform was evenly spread out across the surface using a Drigalski spatula until the chloroform evaporated. Remaining traces of solvent were removed by exposing the dried lipid cake to a vacuum for at least 15 min. Next, 4-5 mL of sodium buffer was added to the petri dish and the solution was left in a humid chamber in the dark at RT for at least 1 h. GVs were detached by gently tapping against the dish and pipetting up and down with a cut pipette tip. The GV suspension was stored at 4 °C in the dark.

When GVs were to be washed and concentrated, the dry lipid film was swollen in caesium buffer or sucrose buffer and, after harvesting, transferred into a falcon tube. The tube was filled up with sodium buffer (isosmotic to the swelling buffers) and centrifuged for at least 30 min at 300 rcf. The supernatant was discarded, and the GV pellet was washed 2-3 more times by adding 900 µl sodium buffer and centrifuging the suspension for 7 min at 300 rcf.

Smaller-sized vesicles were prepared using a mini-extruder (Avanti Research) and polycarbonate membranes with the desired pore size (typically 200 nm; Whatman). GV preparations were pushed back and forth through the membrane 21 times.

Lipids in vesicle preparations were quantified using a colourimetric phospholipid quantification assay kit (Sigma Aldrich CS0001) following the manufacturer’s instructions. The absorbance was measured at 570 nm on a Clariostar plus microplate reader (BMG Labtech).

Vesicles used in the lipid mixture screens (Fig. 2 e,f) were swollen in sucrose buffer at 60 °C to account for differences in lipid charge and melting temperature and washed as described above. Lipid quantity was estimated by measuring the lipid content in sample A (DOPC vesicles) using the colourimetric assay mentioned above. As this assay can only detect choline-containing lipids, the lipid quantity in the remaining samples was estimated based on the fluorescence of Atto643-DOPE species included at equal molar percentages in all lipid mixtures. For this, vesicle preparations were diluted 1:10 and 10 µl aliquots were mixed with 90 µl Triton-X100 (2%) solution in a black 96 well plate. The plate was incubated at 60 °C for 1 h to disintegrate the vesicles and the fluorescence of Atto643-DOPE was measured in a plate reader. The lipid quantity in the other vesicle preparations was finally calculated by normalising the fluorescence readouts in respect to that of sample A, and multiplying the obtained ratio by the colourimetric quantification result for sample A. Images of phase-separated vesicles were obtained using a ThermoFisher EVOS M7000 imaging system.

### Vesicle binding study

Microscope slides and coverslips were washed with ddH_2_O and incubated in denatured BSA blocking buffer (10 mM TRIS, 150 mM NaCl, 50 µM BSA, pH 8) overnight as described previously^33^. Next, the glassware was thoroughly washed with ddH_2_O to remove residual blocking buffer and dried at 50 °C for 1 h. 6 µl of GVs (99.95% DOPC, 0.05% DOPE-Atto488; swollen in imaging buffer A) was mixed with origami triangles (+/- cholesterol modified strands; fluorescently labelled with Atto643 modified oligonucleotides; final origami concentration: 7 nM) and topped up to 40 µl with imaging buffer B. 300 mM NaCl was added to samples testing the effect of elevated sodium on the binding behaviour of origami. The samples were pipetted into an imaging chamber formed by sandwiching a silicon isolator (Grace Bio-Labs) between passivated coverslips and microscope slides. The samples were imaged using a Leica TCS SP5 confocal microscope with a Leica HCX PL APO CS 63x/1.40-0.60 oil immersion objective. Intensity profiles were obtained using Fiji^34^.

### Budding assays

Vesicles were prepared as described above using 99.0 – 100% DOPC with the remainder being fluorescently labelled DOPE if visualisation by fluorescence was desired. Origami triangles were hybridised to cholesterol-modified oligonucleotides by adding them at a 1.5 x excess with respect to the total amount of compatible linker handles on the origami. The hybridisation reaction was incubated at RT for at least 30 min. For outward budding reactions, 24 µl of untreated GV suspension was mixed with origami triangles with linker handles on the shell-inward face at a final concentration of approx. 2 nM origami. In experiments where lipid quantities in GV preparations were known, 1000 pmol lipid of GVs prepared in caesium buffer, or 350 pmol of GVs prepared in sucrose buffer was mixed with the origami (4.5 µl at 15 nM). The sample was topped up to 30 µl with sodium buffer, and the MgCl_2_ concentration was adjusted to 65 mM by adding 1 M MgCl_2_ solution (hypertonic) or to 60 mM by adding isotonic assembly buffer. Budding reactions were incubated at 37 °C for up to 3 d.

For the tonicity screen, undiluted, chol-hybridised triangles were mixed with GVs and diluted using water and glycine buffer. The samples were incubated at RT for 2 h for triangles to bind to the vesicles and for vesicles to adjust to the altered osmotic conditions. Finally, MgCl_2_ solution was added to trigger assembly and samples were incubated at 37 °C overnight. For detailed sample compositions, refer to Supplementary Table 25.

The budding kinetics were obtained by mixing samples with gel loading dye (50% Ficoll400, 20 mM MgCl_2_, Orange G) and freezing them in liquid nitrogen at the indicated time points. Once all time points were taken, the samples were thawed, mixed and used for agarose gel electrophoresis.

For inward budding reactions, origami triangles (3 µl at 20 nM) with linker handles on the shell-outward face were mixed with LVs (1500 pmol lipids; approx. 200 nm diameter) and incubated for up to 3 d.

For bivesicular DNA shells (VCDCVs), triangles with orthogonal linker handles on the top and bottom face were used, and inward and outward budding reactions were conducted successively. First, DCVs were produced in an outward budding assay (1200 µl in total) using cholesterol-modified oligonucleotides hybridised to the bottom face linker handles. The sample was then centrifuged at 300 rcf for 5-8 min, and the pellet containing residual GVs was discarded. The supernatant was then ultracentrifuged at 55 krpm (avg. 108 krcf) for 30 min in a Beckman Coulter Optima MAX-TL ultracentrifuge equipped with a TLA-110 fixed angle rotor. 80-90% of the supernatant was removed, and the pellet was resuspended in the remaining volume.

Meanwhile, LVs (200 nm diameter) were mixed with a second chol-oligo complementary to the linker handles on the top face of the triangles. These oligonucleotides were added at a 50x excess with respect to the total amount of complementary linker handles on the origami, and the solution was incubated at RT for 15-30 min. Then, the magnesium concentration in the LV solution was adjusted to 60 mM MgCl_2_ by adding isotonic assembly buffer. Finally, LVs (1500 pmol lipids) were mixed with the DCV concentrate (to a final concentration of 1-5 nM DNA), and the samples were incubated at 37 °C for 1-3 d.

### Budding using biotin-NeutrAvidin

For budding reactions using biotin-NeutrAvidin interactions to anchor triangles to the membranes, GVs containing 3% biotinylated DOPE, 1% fluorescently labelled DOPE and 96% DOPC were prepared. Origami triangles were folded with linker handles (Supplementary Table 20) complementary to biotinylated oligonucleotides (Supplementary Tables 3) and linker handles for purification using magnetic beads (Supplementary Table 4). Biotinylated oligonucleotides were already included in the folding reaction to remove unbound strands and excess staples by gel extraction. The triangles were then mixed with a 500-fold excess of NeutrAvidin and incubated for 1 h at 37 °C on an orbital shaker. Next, the sample was transferred into a tube previously passivated with denatured BSA blocking buffer and mixed with magnetic beads (Dynabeads M-270 Streptavidin, Invitrogen) coated with dual-biotinylated DNA linker handles (Supplementary Table 4) complementary to the linkers on the origami. The sample was incubated for 1 h at RT on a rotary shaker, and the beads now carrying the origami triangles were washed 3-4 times with sodium buffer. Finally, an invader strand complementary to the full length of the purification linker handles (Supplementary Table 4) was added in great excess, and the solution was incubated at 37 °C for 1 h to uncouple the origami from the magnetic beads by strand displacement. The NeutrAvidin bearing triangles were then mixed with biotinylated GVs, and the MgCl_2_ concentration was adjusted to 60 mM by adding isotonic assembly buffer. The sample was incubated at 37 °C for 2.5 d.

### Negative stain transmission electron microscopy

Sample solutions were incubated on glow-discharged copper grids (400 mesh) with a carbon support film (Science Services) for 4-8 minutes, depending on the sample concentration. The solution was then blotted off using filter paper (Whatman). Then, the grid was washed once using stain solution (2% aqueous uranyl formate + 25 mM NaOH), followed by incubating another stain droplet on the grid for 30 s. Excess stain was blotted off using filter paper, and the grid was air-dried before imaging.

Grids were typically imaged at x26-30k magnification using SerialEM with an FEI Tecnai T12 electron microscope operated at 120kV and a Tietz TEMCAM-F416 camera. Image contrast was auto-levelled using Fiji.

For tomography, one-directional tilt series were acquired from −50° to +50° in steps of 2° using SerialEM. The resulting image stacks were imported into Etomo (IMOD^35^), and images were aligned without fiducials by cross-correlation with cumulative correlation switched on. Tomograms were generated by filtered back-projection (gaussian filter cut-off: 0.35; fall-off: 0.035;). For visualisation, the z-stacks thus generated were imported into Fiji and z-projected by summing up z-slices corresponding to the top, middle and bottom sections of the respective particle of interest.

### Cryogenic electron microscopy

Sample grids for cryoEM were prepared using an automated vitrification system (FEI Vitrobot Mark V). First, grids (Quantifoil R1.2/1.3, Cu 200 mesh, 100 holey carbon films, 2 nm carbon support film; for images of DCVs: Ted Pella, Cu 400 mesh, lacey carbon, <3 nm carbon support film) were glow-discharged and then inserted into the Vitrobot. The conditions inside the device’s environmental chamber were kept at either 4 °C or 22 °C and 100% relative humidity. Samples were typically incubated on the grids inside the chamber for 5 min and plunge-frozen in liquid ethane (blot force: −1, 0 or 25; blot time: 2-3 s; blot total: 1; drain time: 0 s;). The grids were imaged with a spherical-aberration (Cs)-corrected Titan Krios G2 electron microscope (Thermo Fisher) operated at 300kV and equipped with a Falcon III 4k direct electron detector (Thermo Fisher). Samples were imaged at x29k magnification and defocus values between −1 and −4 µm. Images were auto-levelled in Fiji.

### Agarose gel electrophoresis and image processing

Agarose gels were cast by dissolving UltraPure agarose (Invitrogen) in running buffer (0.5 x TBE, Carl Roth) supplemented with MgCl_2_ and optionally ethidium bromide (0.025%, Carl Roth). Samples were run for 2 h at 90 V and then scanned using a Typhoon FLA 9500 laser scanner (GE Healthcare) at a pixel size of 50 µm. Gels containing only monomeric origami structures were run with 5.5 mM MgCl_2_ in the gel and running buffer and an agarose concentration of 1.5%. To resolve and preserve assembled shell structures, the gel and running buffer were supplemented with 20 mM MgCl_2_ and the agarose concentration was reduced to 0.8%. Due to the elevated salt concentration in these gels, the gel boxes were cooled in a water-ice bath, and the running buffer was exchanged every 45 min to replenish lost magnesium. For visualisation, brightness/contrast of the area underneath the gel pockets was auto-levelled in Adobe Photoshop and Fiji, intentionally oversaturating the gel pockets in the process.

Gels containing both fluorescent DNA and lipids were scanned using appropriate excitation wavelengths and emission filters. A merged-colour image was created in Fiji, setting DNA signal grey and lipid signal magenta.

## Acknowledgements

This manuscript has been edited in part with the assistance of the large language models GPT3.5 (openAI) and Perplexity AI. We thank Andreas Bausch and Friedrich Simmel for technical assistance, Björn Högberg for discussions, and Volodymyr Mykhailiuk for help with illustrations. We acknowledge financial support received from the European Research Council Advanced Grant awarded to H.D. (grant agreement 101018465) and the Deutsche Forschungsgemeinschaft for the Gottfried Wilhelm Leibniz Program grants provided to H.D. This project has received funding from the European Union’s Horizon 2020 research and innovation programme under the Marie Skłodowska-Curie grant agreement No 765703.

## Author contributions

The research was performed by M.T.P., supervised by H.D. The manuscript was written by M.T.P. and H.D.

## Data availability

All data are available in the main text or supplementary materials.

## Supplementary Materials

**Supplementary Figure 1.**
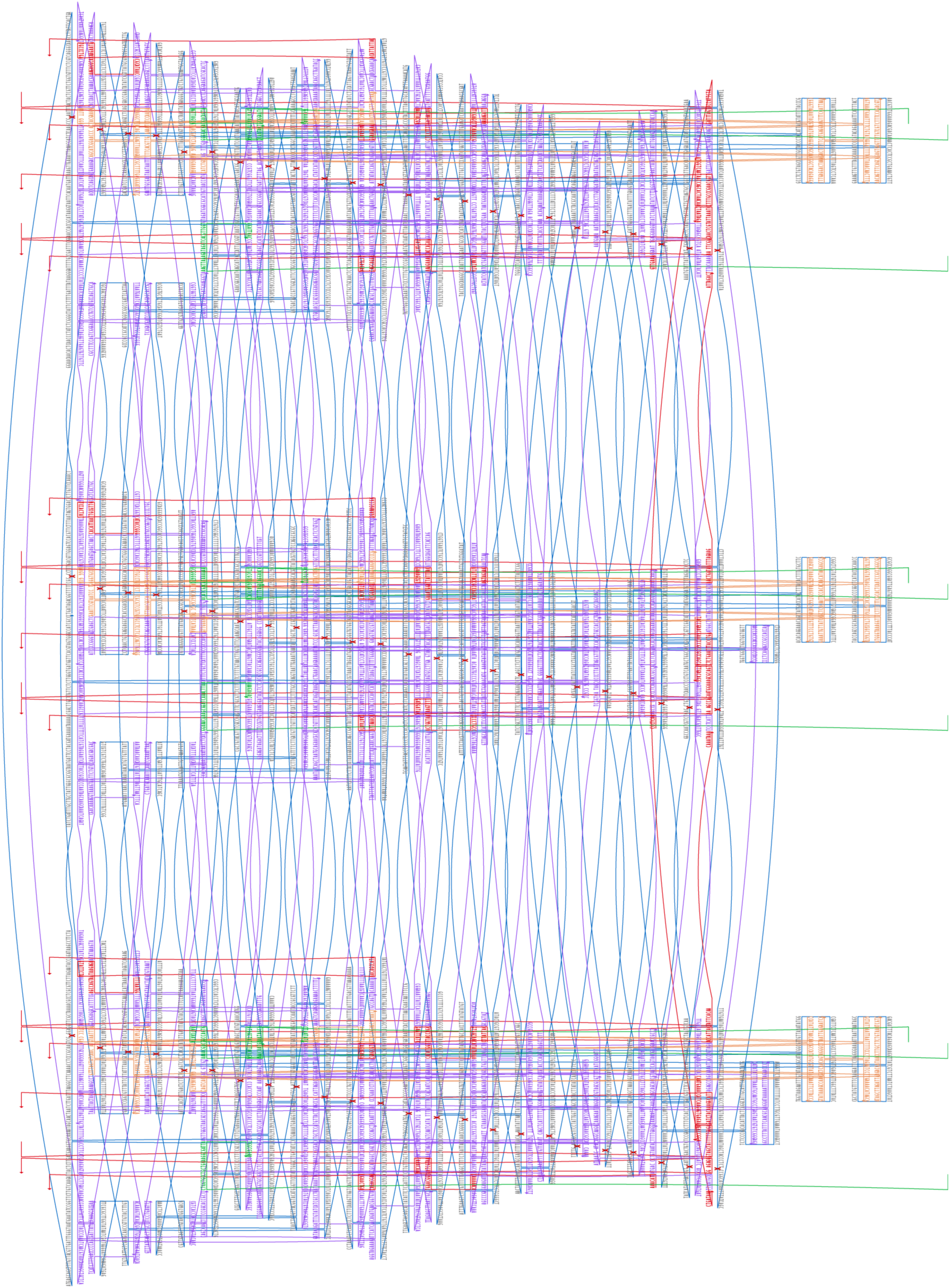
Triangle design map. Exemplary linker handles extending from the origami surface are shown in red (shell-outer face) and green (shell-inner face). The three main domains represent the three sides comprising the triangle. Protrusions used for triangle assembly are highlighted in orange, the shape-complementary recesses are visible as holes in the design.

**Supplementary Figure 2.**
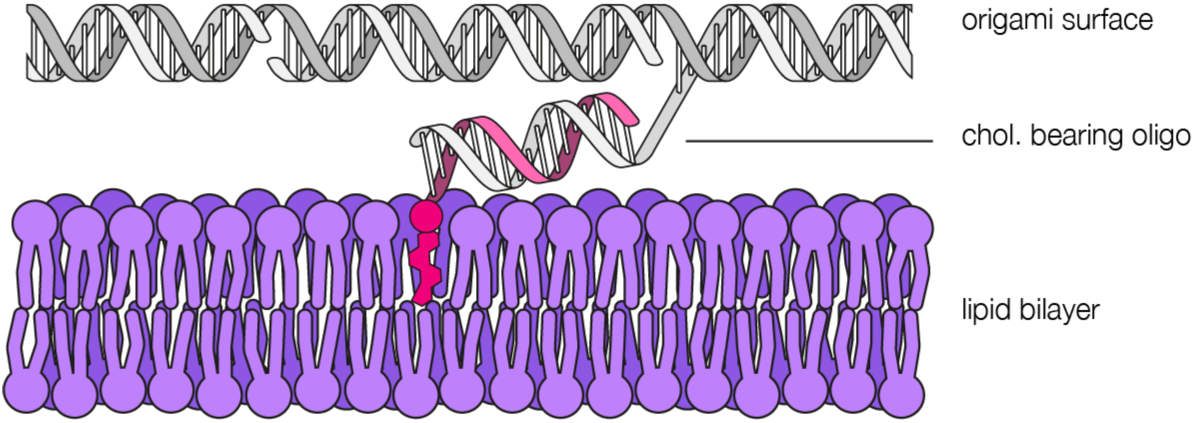
Attachment of origami triangles to vesicles via cholesterol-bearing oligonucleotides. Linker handles extending from the origami surface are hybridised to cholesterol-bearing oligonucleotides with complementary sequences. Cholesterol acts as a lipid membrane anchor to tether the triangles to lipid vesicles.

**Supplementary Figure 3.**
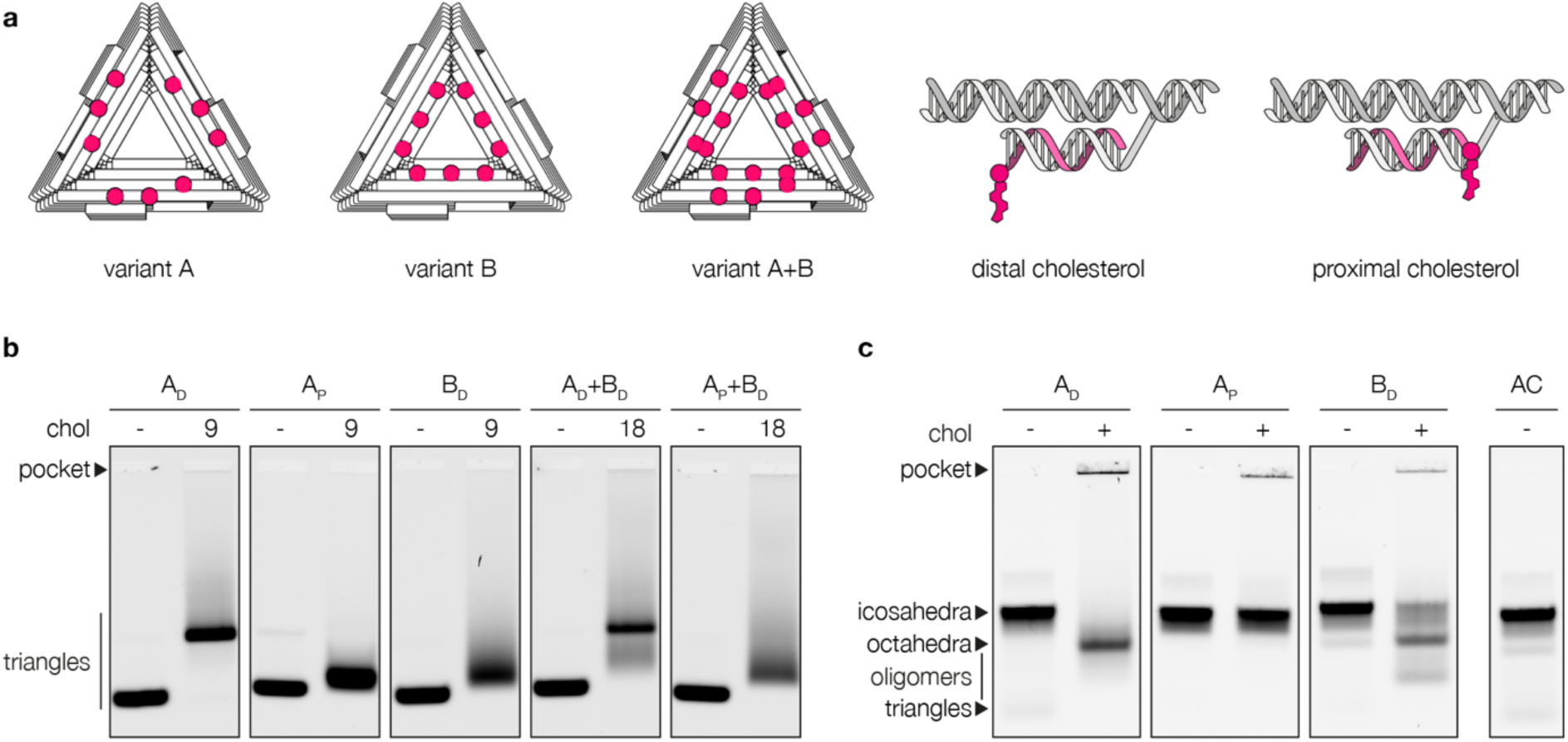
The influence of cholesterol position and orientation on origami triangles. **a,** Linker handle positions on three different triangle variants, and scheme of distal and proximal cholesterol configurations. The upper, white helix represents the triangle surface (side view), the pink strand represents the cholesterol-bearing oligo (chol-oligo). Cholesterol may be oriented facing away (distal) or towards (proximal) the origami surface. **b,** The overall reach of chol-oligos determines the migration behaviour of origami triangles. When cholesterols are positioned closer to the outer edge and the stacking contacts in a distal configuration (A_D_), bands in an agarose gel appear shifted and smeared compared to the same triangles not hybridised to chol-oligos. This effect could be reduced by changing the orientation of the cholesterol to a proximal configuration with respect to the origami, such that the reach of the cholesterol is minimised (A_P_). When chol-oligos are positioned closer to the centre of the structure, and thus further away from the stacking contacts, the migration distance of these triangles is comparable to that of sample A_P_, albeit with more smearing. Combining both variants to double the cholesterol count per triangle from 9 to 18 moieties yielded bands appearing as combinations of the individual variants. **c,** The reach of cholesterol influences the assembly of cholesterol-decorated triangles. An assembly control (AC) of triangles lacking protruding linker handles assembles mostly into icosahedral species, as do triangles with protruding linker handles not hybridised to chol-oligos. When chol-oligos are introduced, distal cholesterol (A_D_, B_D_) promotes the formation of primarily octahedral species, whereas proximal cholesterol (A_P_) does not noticeably influence the assembly behaviour. Reducing the reach of individual cholesterols by positioning them closer to the centre of the triangles and choosing proximal configurations reduces unwanted interactions.

**Supplementary Figure 4.**
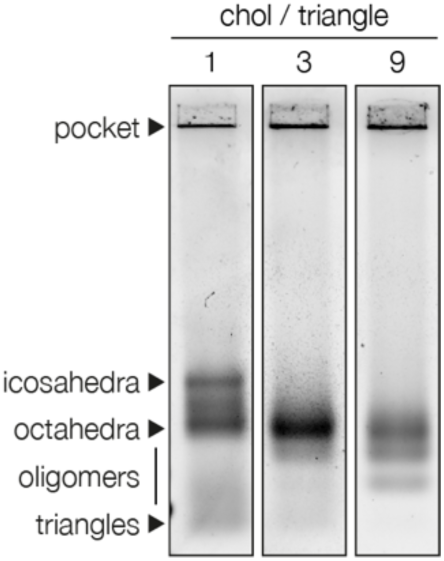
The influence of cholesterol count per triangle in assembly reactions containing GVs. Whilst optimising cholesterol positioning on the triangle can reduce unexpected assembly behaviour, introducing lipid vesicles yet again shifts the assembly towards octahedra, the degree of which increases with the number of cholesterols per triangle.

**Supplementary Figure 5.**
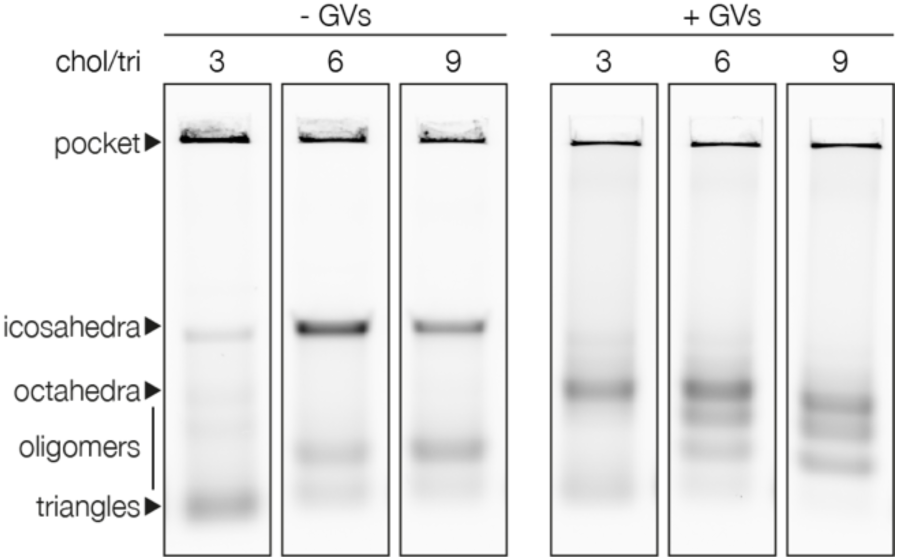
GVs shift the assembly of cholesterol-bearing triangles towards octahedral shells. The assembly behaviour of cholesterol-bearing triangles, tagged with Atto643 for visualisation, depends on the presence or absence of GVs. Without GVs, triangles mainly assemble into icosahedral shells as per their design. With GVs, however, most shells detected by agarose gel electrophoresis are octahedral. The abundance of smaller, intermediate species between closed shells and monomeric triangles increases with the number of cholesterols per triangle and mostly represents aggregated triangles.

**Supplementary Figure 6.**
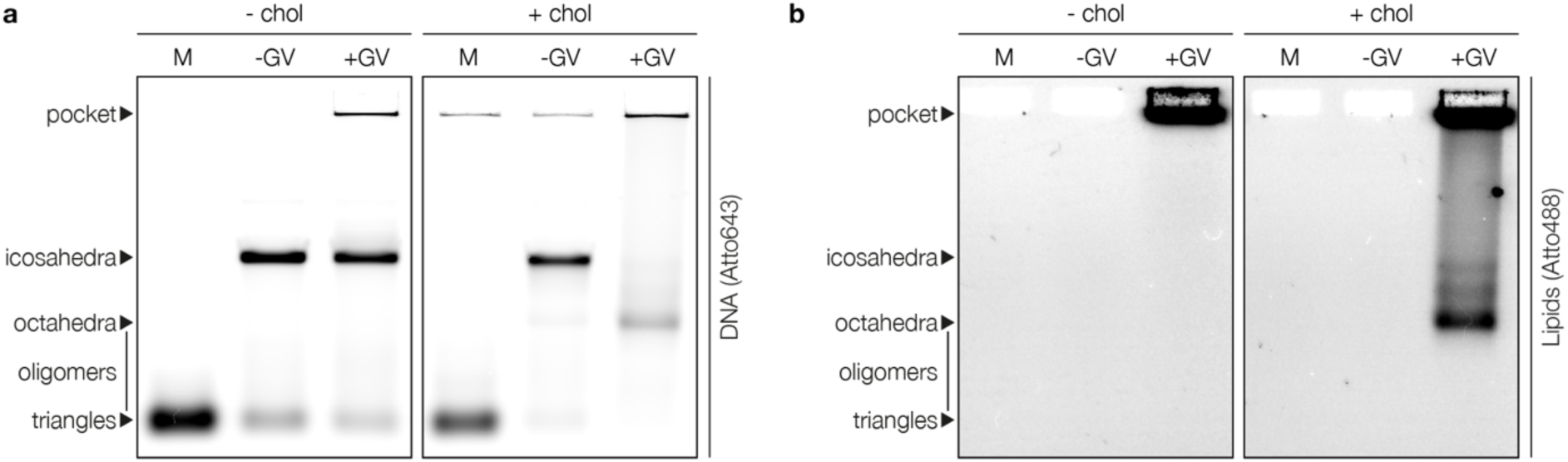
GVs only alter the assembly behaviour of chol-bearing triangles. **a,** Agarose gel of DNA shells made visible by Atto643-bearing oligonucleotides.Triangles assemble predominantly into icosahedral shells even if either GVs or chol-bearing oligonucleotides are present in the sample. However, if both GVs and chol-bearing oligonucleotides are present in a sample at the same time, the assembly of triangles is shifted towards octahedral species. **b,** The gel in a rescanned for visualisation of DOPE-Atto488 species included in the vesicle membranes. Only the sample containing chol-bearing triangles and GVs produced lipid bands coinciding with the DNA shell bands in a, therefore indicating that the octahedral shells found in a are mostly octahedral DNA-shell-coated vesicles. (M: triangle monomers)

**Supplementary Figure 7.**
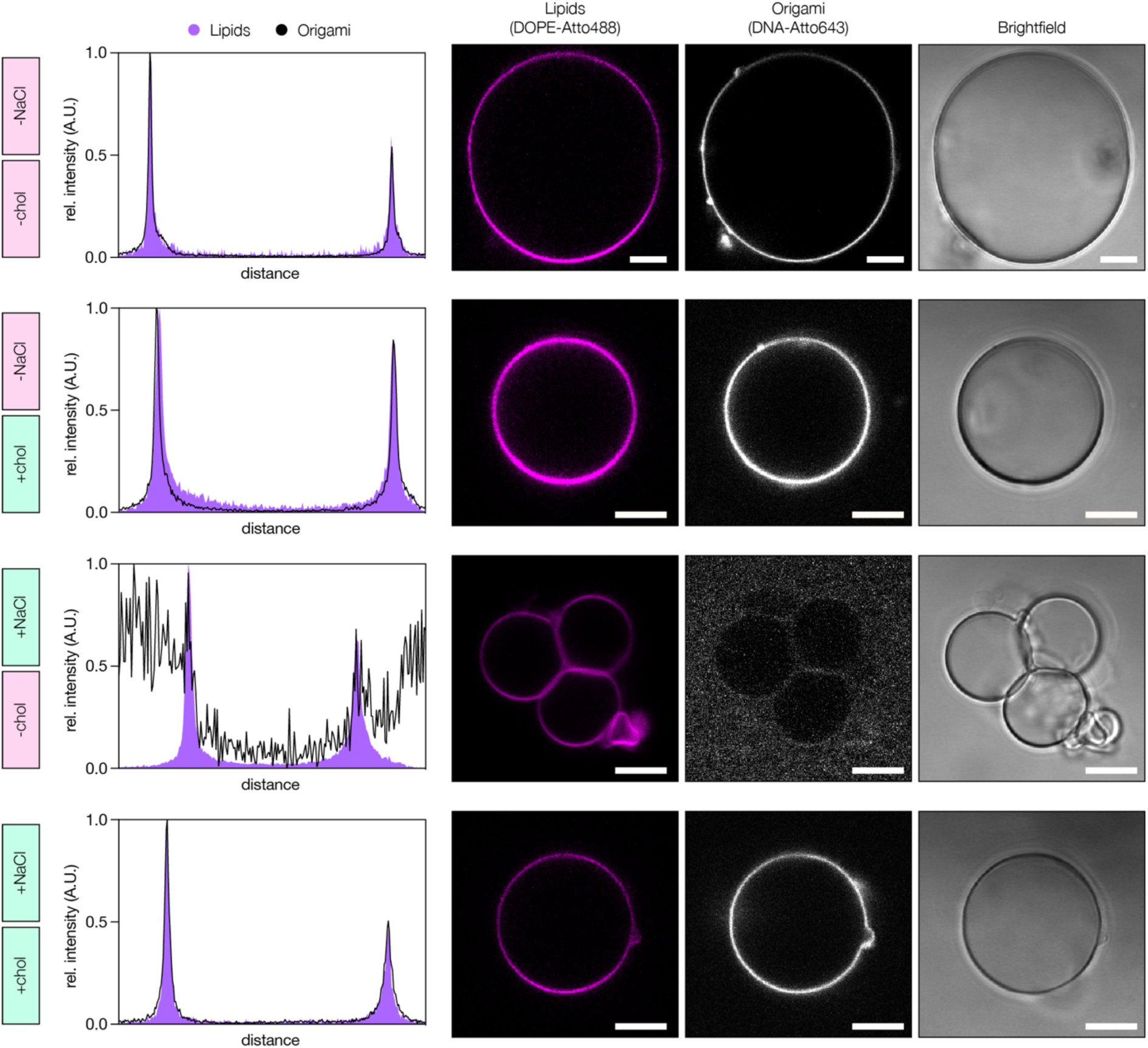
Sodium chloride prevents non-specific adsorption of DNA origami onto giant vesicles. Cross-sectional intensity profiles and channel-separated confocal microscopy images of giant vesicles mixed with DNA origami triangles. Conditions: +/- cholesterol-modified oligonucleotides (chol) and +/- 300 mM NaCl, as indicated. At low-salt conditions, triangles adsorb onto lipid vesicles even if not hybridised to chol-oligos. By adding 300 mM NaCl, non-specific origami-vesicle association is suppressed without interfering with cholesterol-mediated association. Scale bars: 10 µm

**Supplementary Figure 8.**
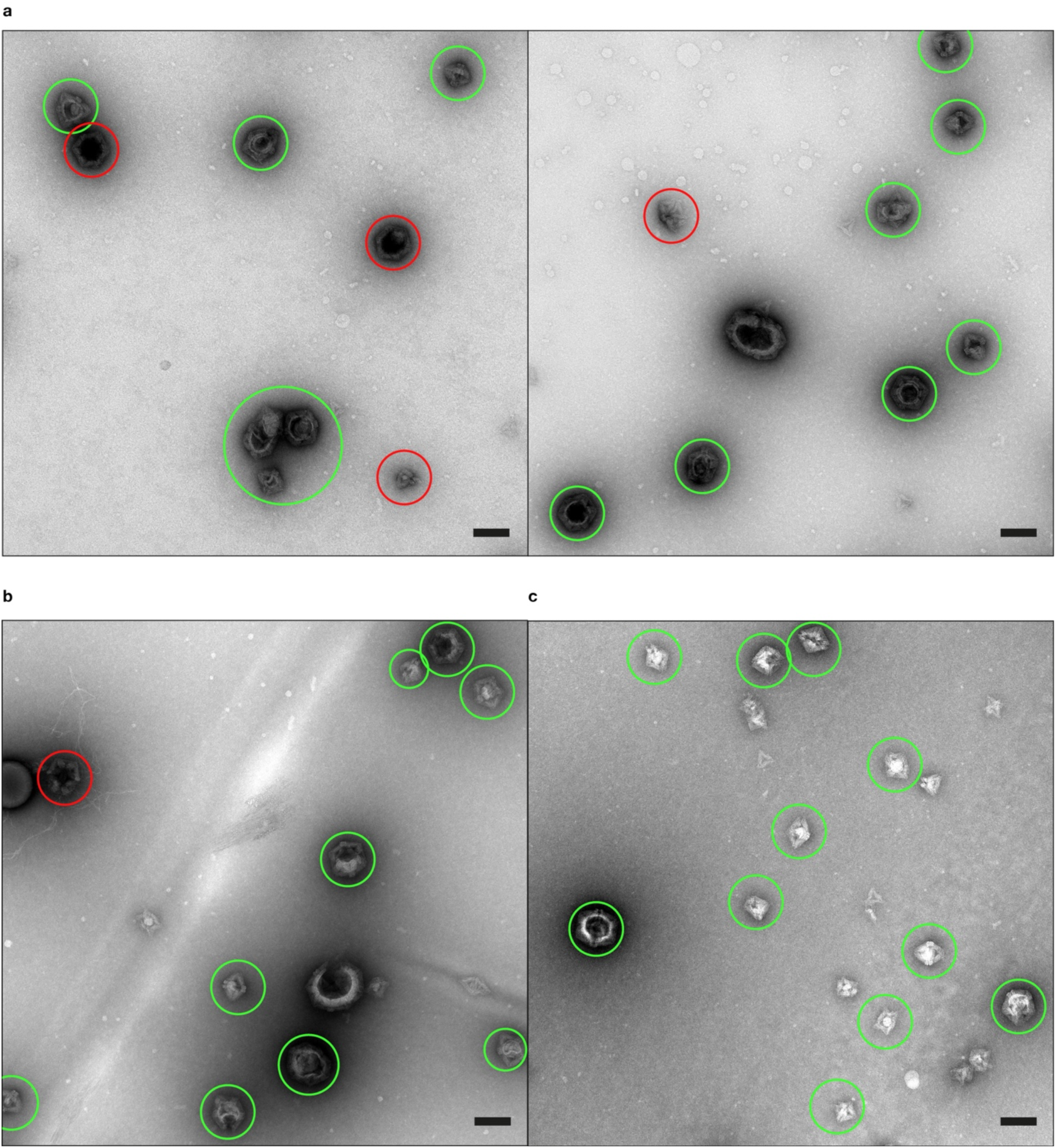
DCVs obtained from triangles with 3, 6 or 9 cholesterols. Full fields-of-view of DCV samples imaged using negative stain TEM. DCVs are encircled green and empty shells are encircled red. Particles with ambiguous appearance, DCVs with irregular shapes (e.g. particles larger than octa- and icosahedral particles), and mono- or oligomeric triangles have not been marked. The micrographs were obtained under the same conditions from triangles carrying either **a,** 3, **b,** 6, or **c,** 9 cholesterol moieties. Under optimised conditions, DCVs make up the largest particle fraction. Scale bars: 100 nm.

**Supplementary Figure 9.**
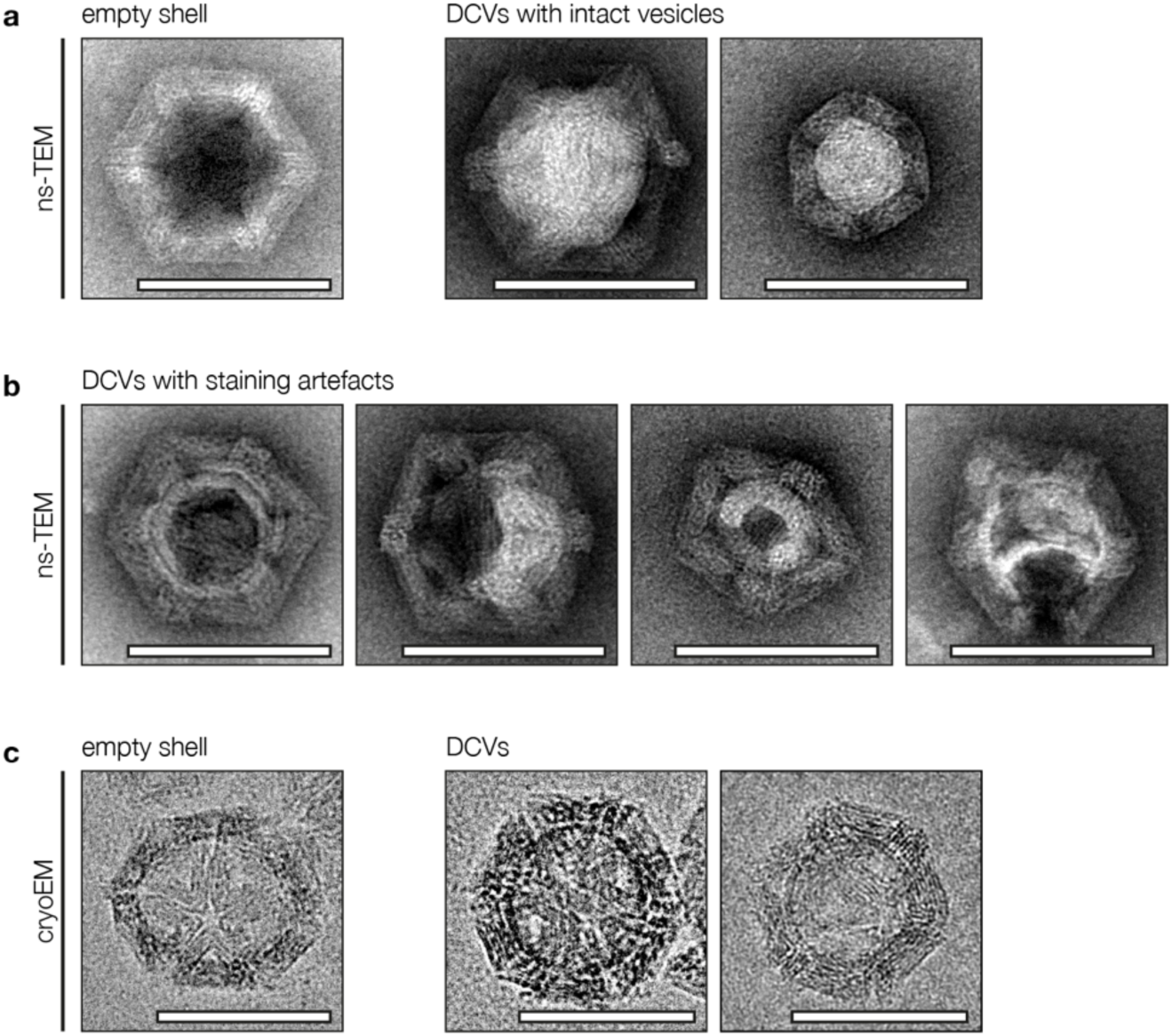
Appearance of DCVs in TEM micrographs. **a,** Uranium stain for negative stain TEM (ns-TEM) increases the contrast of biological samples, allowing for easy differentiation between different shell species. Empty shells accumulate stain within their cavity, rendering it black, while lipid vesicles within DCVs displaces the stain, rendering the cavity white. **b,** Staining and subsequent drying of sample grids commonly alters the appearance of lipid vesicles. Consequently, the vesicles within DCVs commonly appear distorted, incomplete or broken in ns-TEM. **c,** Cryogenic electron microscopy (cryoEM) allows imaging of DCVs in their native state without sample staining or drying steps. DCVs can be distinguished from empty shells by a dark ring lining the inner cavity of the DCV, representing the lipid vesicle. The artefacts shown in b are absent in cryoEM micrographs. All scale bars: 100 nm

**Supplementary Figure 10.**
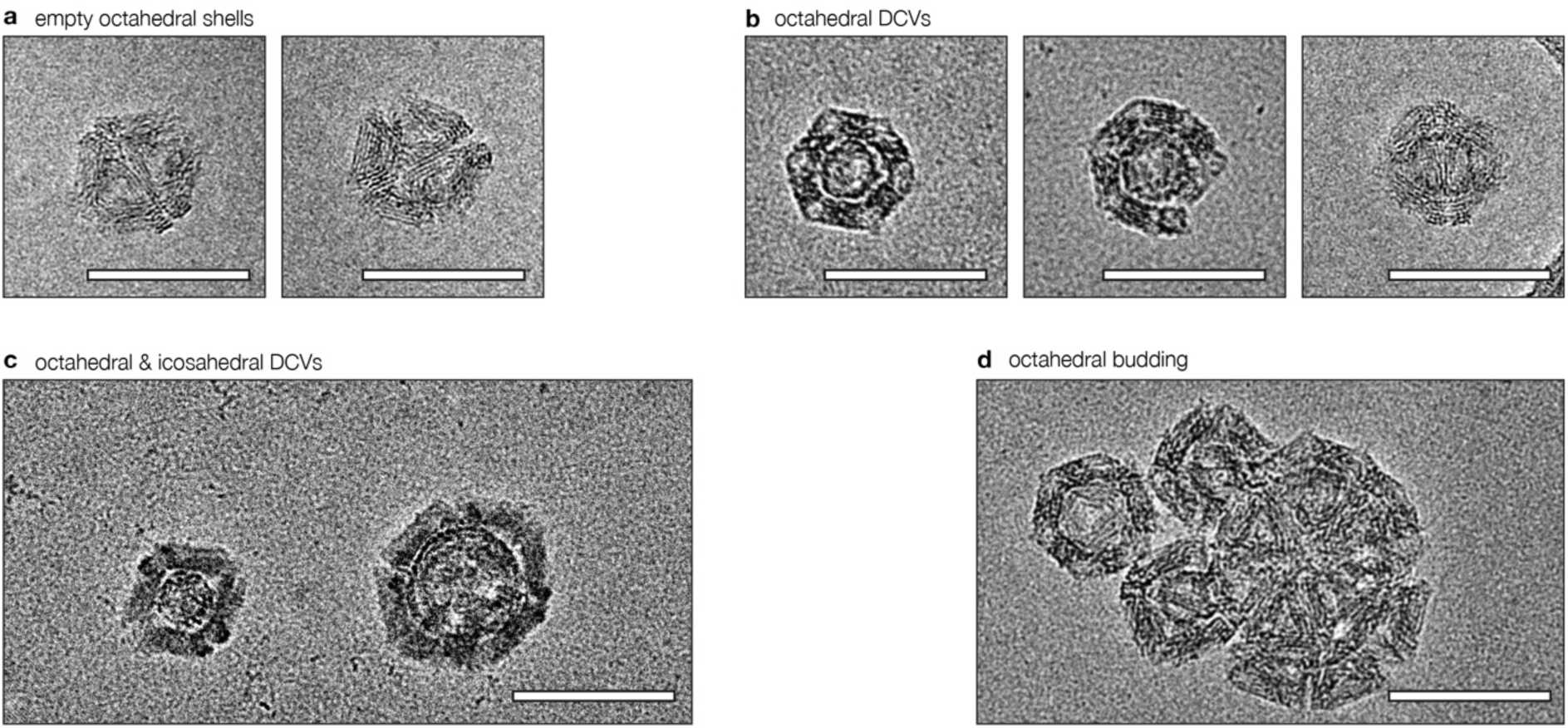
CryoEM images of octahedral DCVs. **a,** Appearance of empty octahedral shells. **b,** Octahedral DCVs. Differences in appearance are due to different orientations of the particles. **c,** Size comparison between an octahedral (left) and icosahedral (right) DCV. As icosahedral particles are composed of 20 triangles (versus 8 in octahedra), the inner cavity and the overall size of the particle are both larger than in octahedral species. **d,** Budding of two octahedral DCVs. To the left of the two triangle-covered buds is an empty octahedral shell. All scale bars: 100 nm.

**Supplementary Figure 11.**
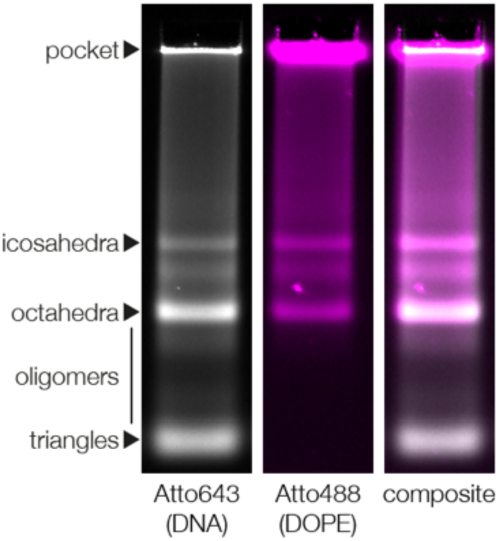
Identification of DCVs in agarose gel electrophoresis by colocalisation analysis. The relative efficiency of DCV formation can be analysed by agarose gel electrophoresis using fluorescently labelled DNA oligos attached to the origami structure or ethidium bromide, and a small fraction of fluorescently labelled DOPE species in the lipid mixture comprising the vesicles. Whereas multiple bands can typically be detected in the DNA channel, representing triangles in monomeric, oligomeric or fully assembled states, lipid signal was only seen at the height of assembled shells. The absence of lipid signal colocalised with monomeric triangles suggests that lipids predominantly enter the gel as part of a DCV. The gel pockets contain origami-coated GVs, which were oversaturated to better visualise the much fainter bands below.

**Supplementary Figure 12.**
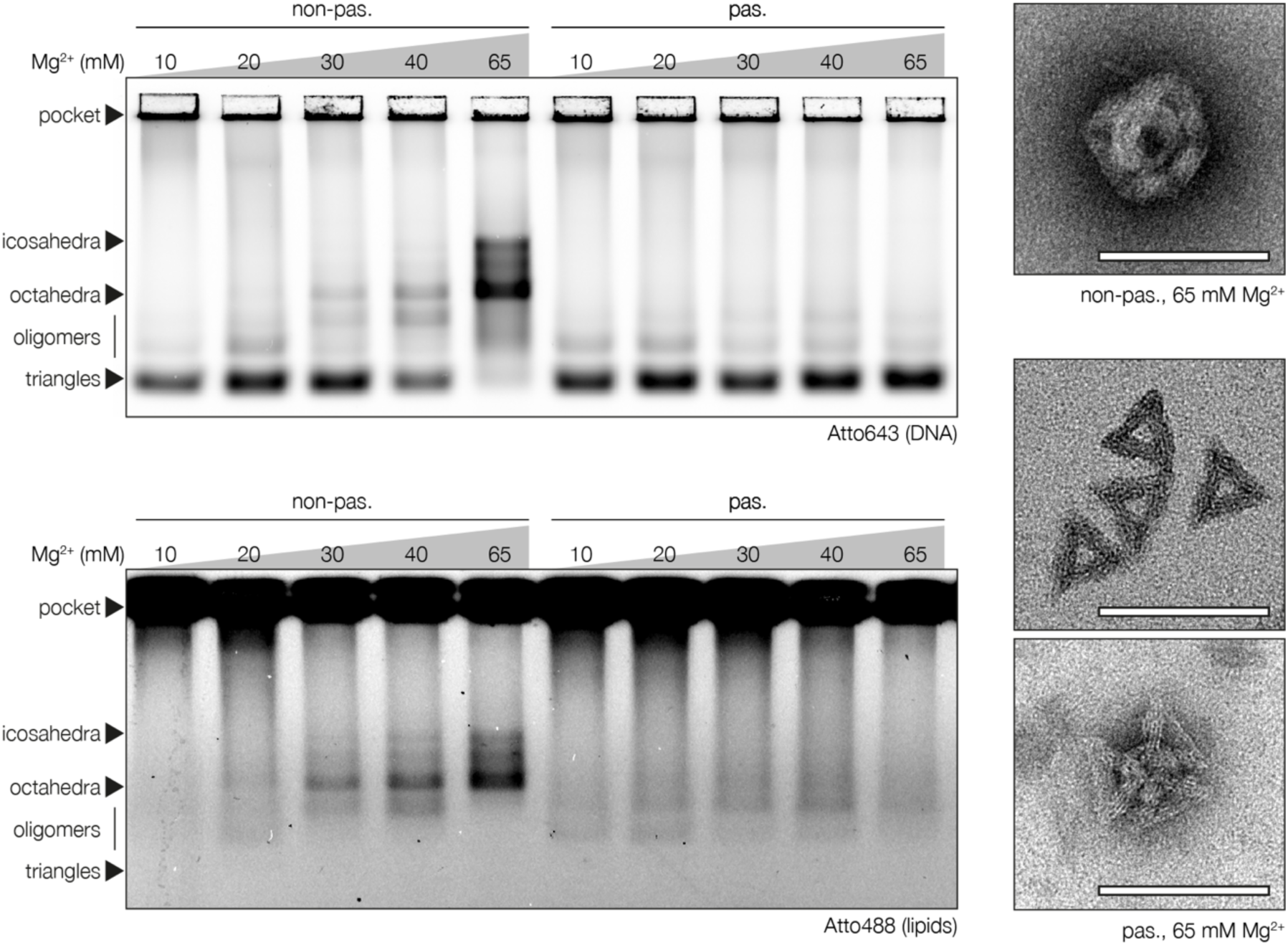
Triangle assembly is the driving force behind DCV formation. The importance of shell assembly for DCV formation was studied by incubating triangles at various MgCl_2_ concentrations at (65 mM) or below (10-40 mM) the typical threshold concentration for shell assembly. For visualisation by agarose gel electrophoresis, the triangles were hybridised to Atto643 bearing oligonucleotides, and the GVs contained a fraction of Atto488 labelled DOPE. Two triangle species were prepared: A non-passivated version with blunt-ended stacking contacts and a passivated version where staple oligonucleotides at the base stacking contacts were extended by additional unpaired thymidines. Whereas the non-passivated versions formed DCVs, as seen by the bands in the lipid channel (lower panel) colocalising with the shells in the DNA channel (upper panel), only faint bands below the typical height of DNA shells were found in samples containing the passivated version. TEM analysis (right panels) revealed that while the passivated triangle samples contained mostly monomeric triangles, there was a small number of aggregated triangles held together by small vesicles, likely representing the faint bands. The lack of DCVs in samples containing passivated triangles highlights the importance of shell assembly for DCV formation. Scale bars: 100 nm.

**Supplementary Figure 13.**
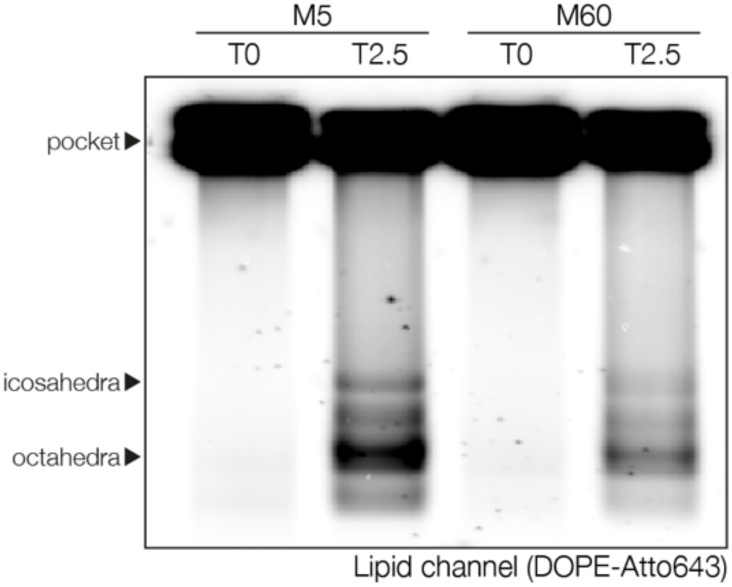
Shell preassembly reduces DCV yields. Cholesterol-bearing origami triangles were incubated without GVs for 4 h at 37 °C in either 5 mM (monomer conditions) or 60 mM (assembly conditions) MgCl_2_ to study the effect partially assembled shells have on DCV yields. Afterwards, GVs (containing a small fraction of DOPE-Atto643) were added, and the MgCl_2_ concentration in the 5 mM controls was increased to 60 mM. Aliquots taken and frozen at t = 0 d were compared to samples incubated at 37 °C for 2.5 d using agarose gel electrophoresis, and DCV yields were analysed by comparing lipid bands. Whereas t = 0 d aliquots looked alike in samples pre-incubated in either 5 mM or 60 mM MgCl_2_, we finally obtained stronger lipid signals in samples initially incubated at 5 mM. We deduce that DCVs form best when monomeric triangles are assembled directly on the membrane. Partially assembled shells may fail to bind GVs efficiently due to steric hindrance, resulting in reduced DCV yields.

**Supplementary Figure 14.**
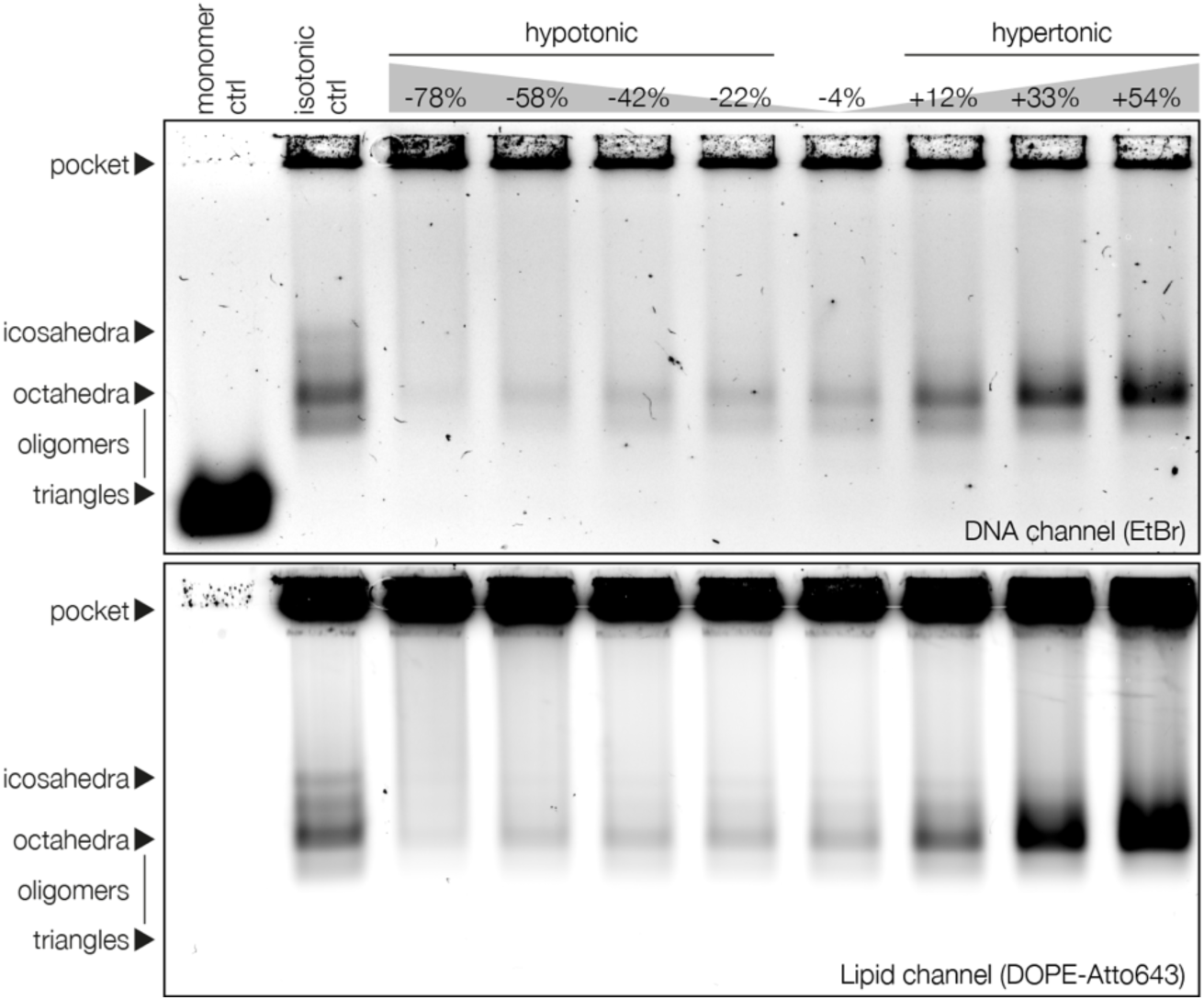
Budding assays at varying tonicities (channel-separated). Agarose gel of budding assays ran at different tonicities and visualised using two fluorophores. The top image shows signal obtained from DNA origami stained by ethidium bromide, while the bands in the bottom image stem from Atto643-labelled DOPE species included in the lipid mixture of the GVs. Budding efficiency strongly correlates with tonicity, working best under hypertonic conditions and coming to a near halt at the most hypotonic condition tested. Refer to Figure 2c for a merged-colour image.

**Supplementary Figure 15.**
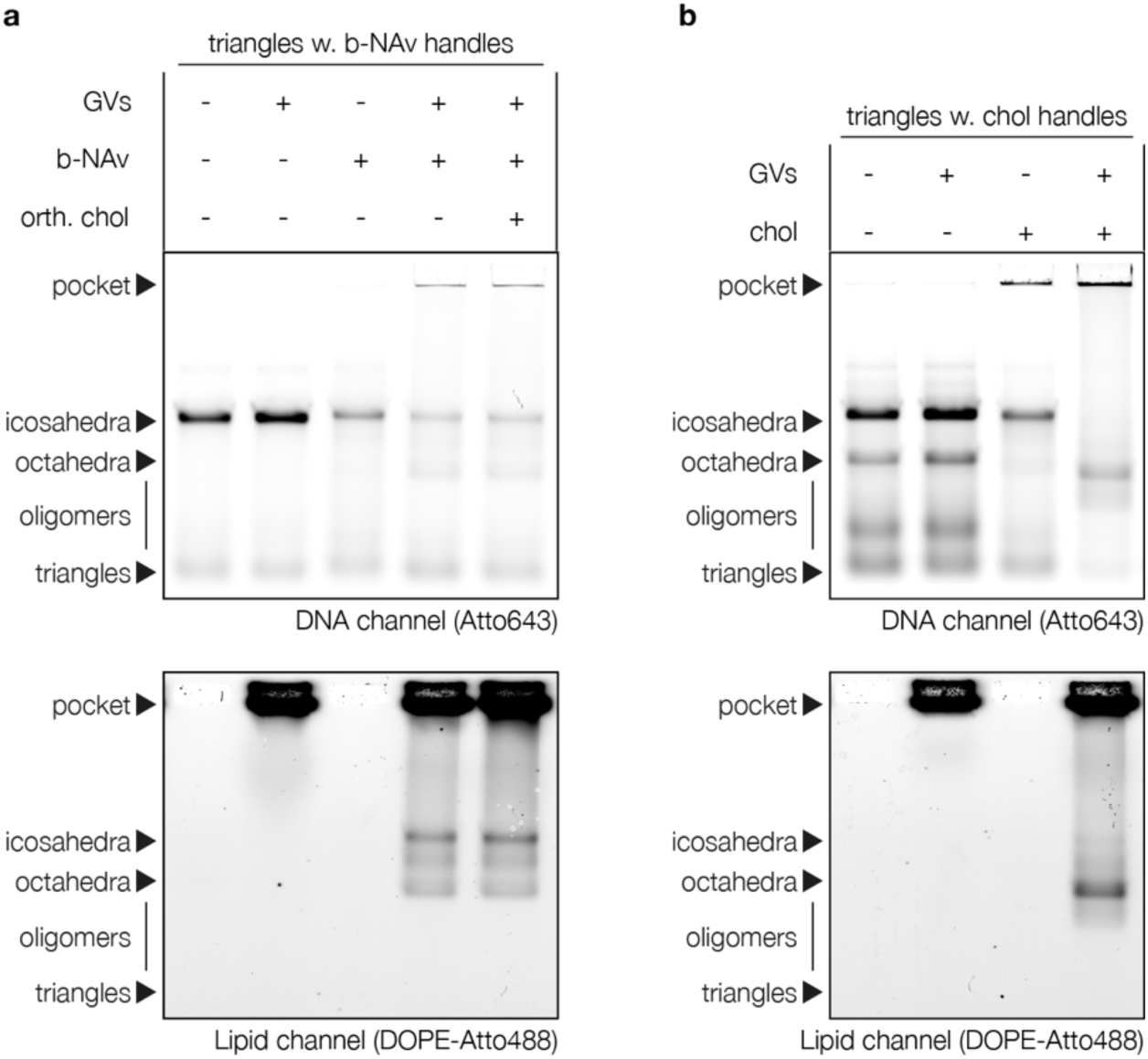
Membrane anchor screen (channel-separated). **a,** Agarose gel of DCVs prepared by using biotin-NeutrAvidin (b-NAv) interactions to connect origami triangles (Atto643, top image) and GVs (DOPE-Atto488, bottom image) containing a fraction of biotinylated DOPE. Linker handles on the triangles were complementary to b-NAv-oligos, but not to chol-oligos. Lipid material remains in the pockets unless triangles are mixed with b-NAv-oligos and GVs. The bands in the lipid channel colocalise with polyhedral shells but not with monomeric triangles, confirming the formation of DCVs. The addition of orthogonal chol-oligos did not improve yields, showing that the insertion of cholesterols into the bilayer is not a requirement. Gel pockets were oversaturated for better band visualisation. Refer to Figure 2d for a merged-colour image. **b,** Agarose gel of DCVs prepared under the same conditions as in a, using triangles with linker handles complementary to chol-oligos.

**Supplementary Figure 16.**
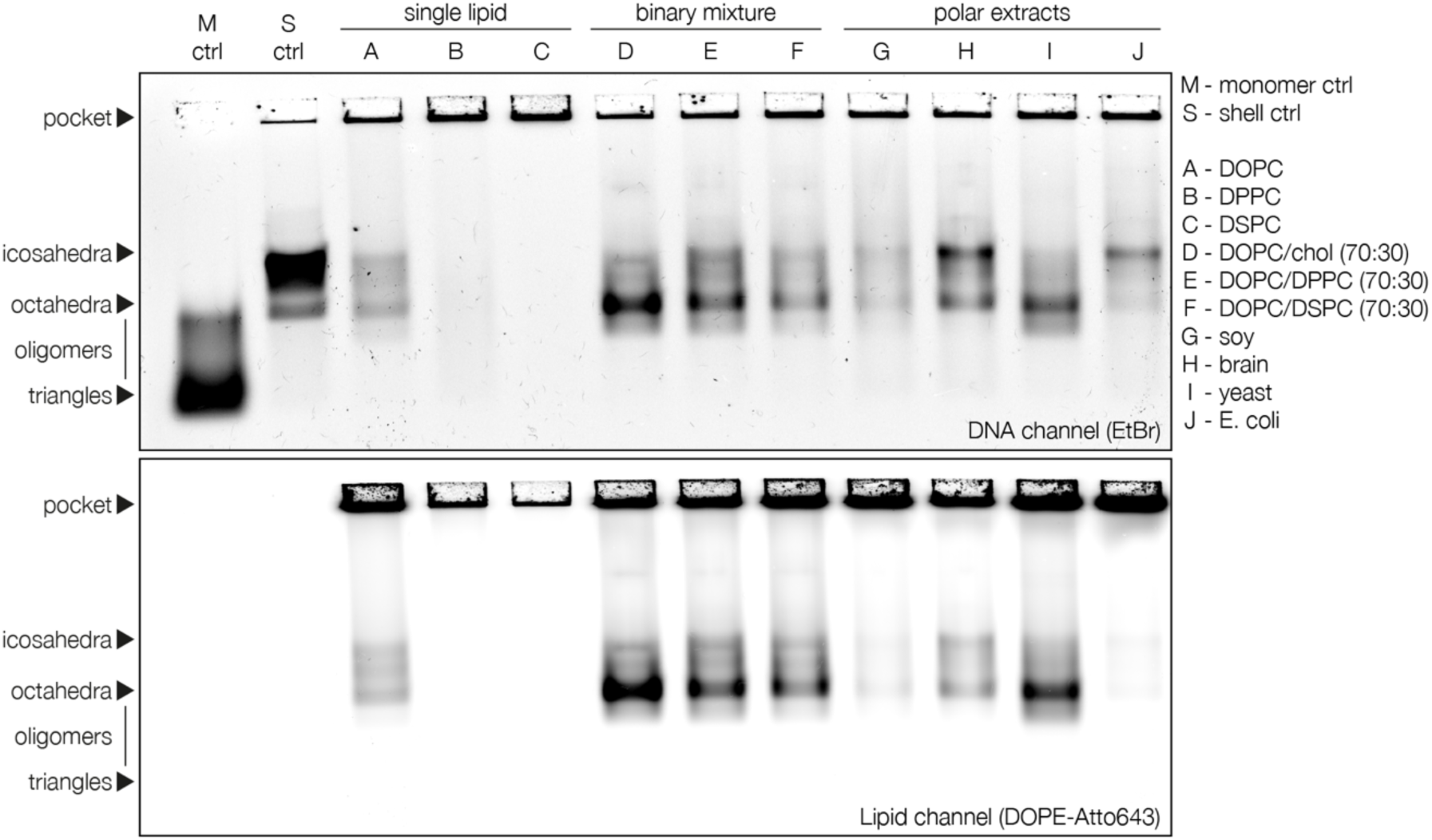
Budding assay using giant vesicles of varying lipid composition (channel-separated). Agarose gel of DCVs obtained from GVs of various lipid compositions. All compositions tested allowed budding of DCVs at varying efficiencies, except for GVs composed entirely of high-melting lipids. Top image: DNA (EtBr); Bottom image: Lipids (DOPE-Atto643); Refer to Figure 2e for a merged-colour image.

**Supplementary Figure 17.**
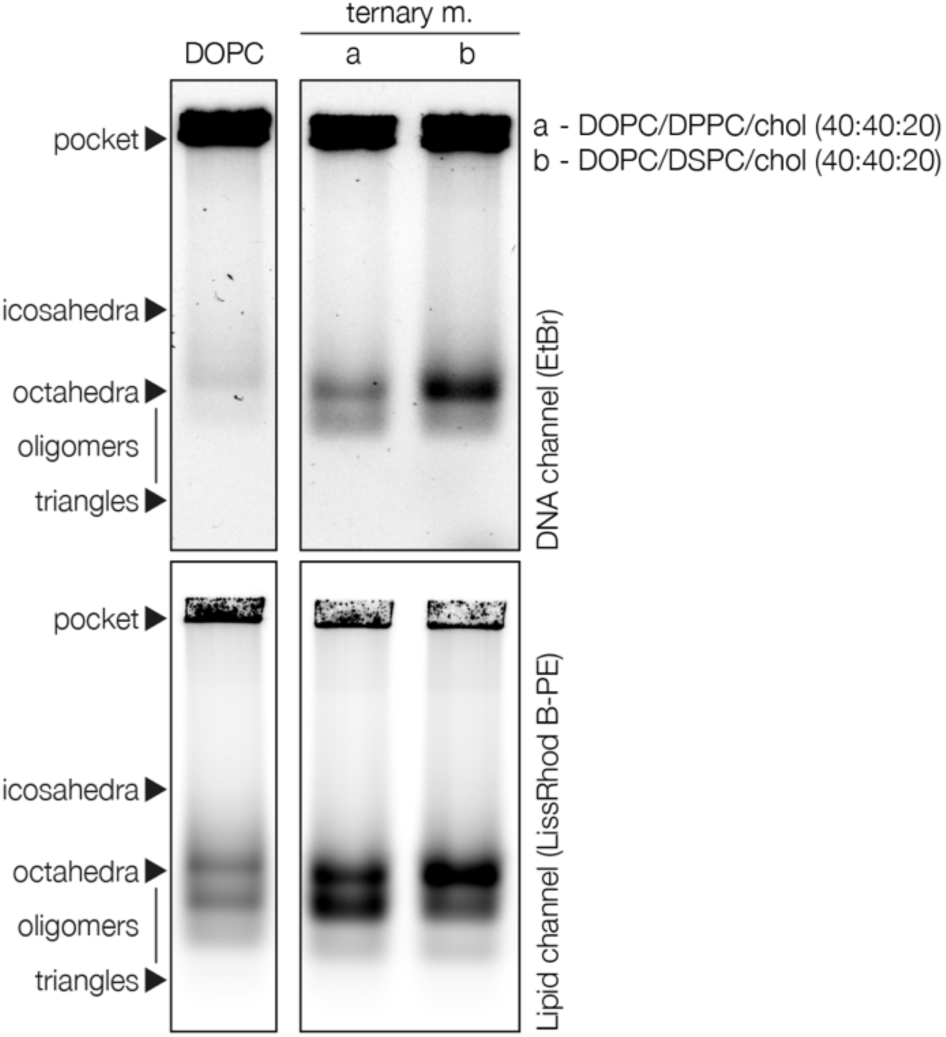
Budding assay from phase-separated vesicles (channel-separated). Agarose gel comparing DCVs obtained from GVs composed of DOPC or phase-separated ternary lipid mixtures. Phase-separated vesicles yielded more DCVs, suggesting an influence of phase boundaries on the budding mechanism. Top images: DNA (EtBr); Bottom images: Lipids (LissRhod-PE); Refer to Figure 2f for a merged-colour image.

**Supplementary Figure 18.**
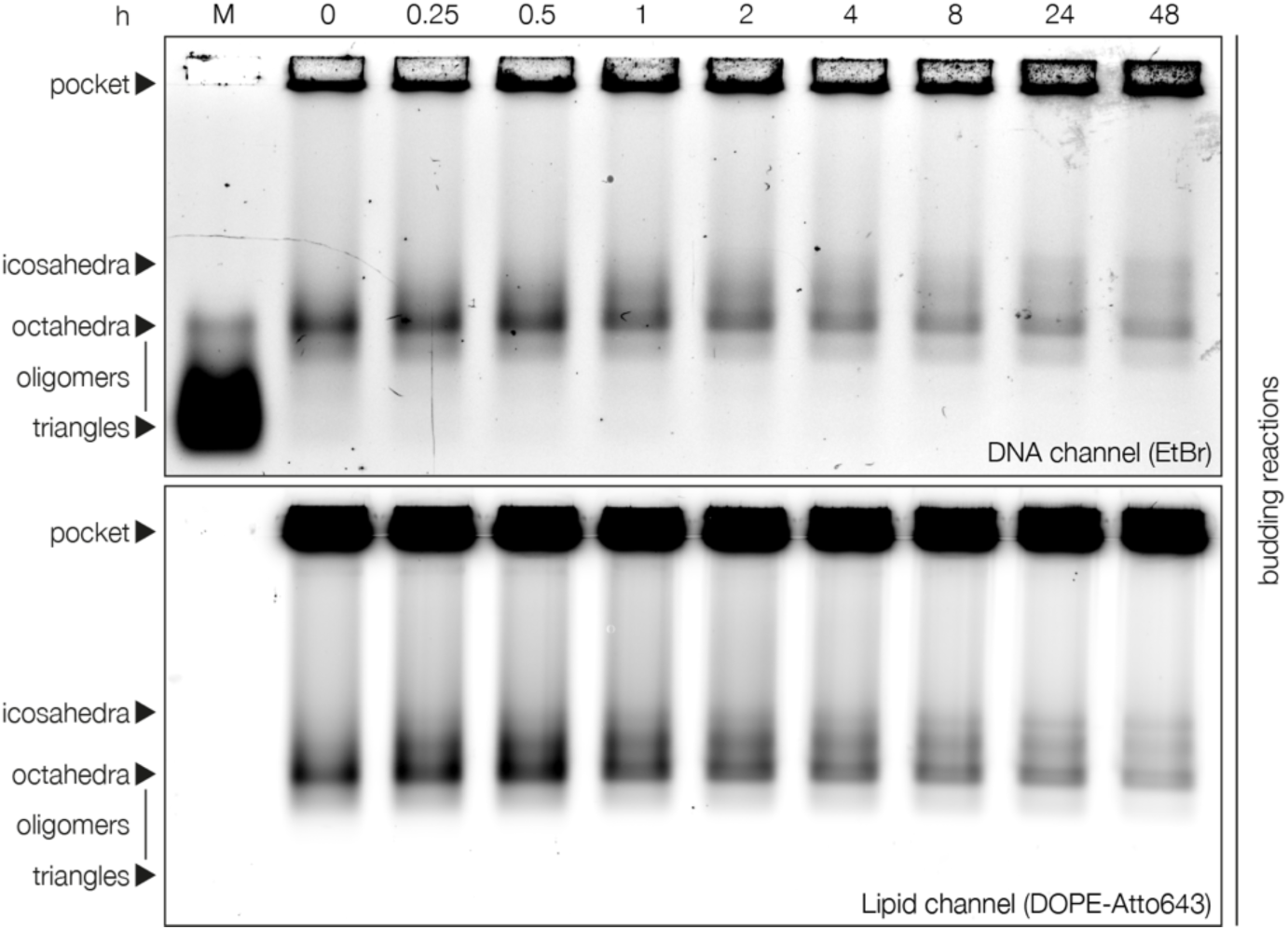
Budding kinetics (channel-separated). DCV budding occurs instantly and yields mostly octahedral species, but at longer incubation times a notable fraction of icosahedral DCVs is formed. Top image: DNA (EtBr); Bottom image: Lipids (LissRhod-PE); Refer to Figure 2g for a merged-colour image.

**Supplementary Table 1.**
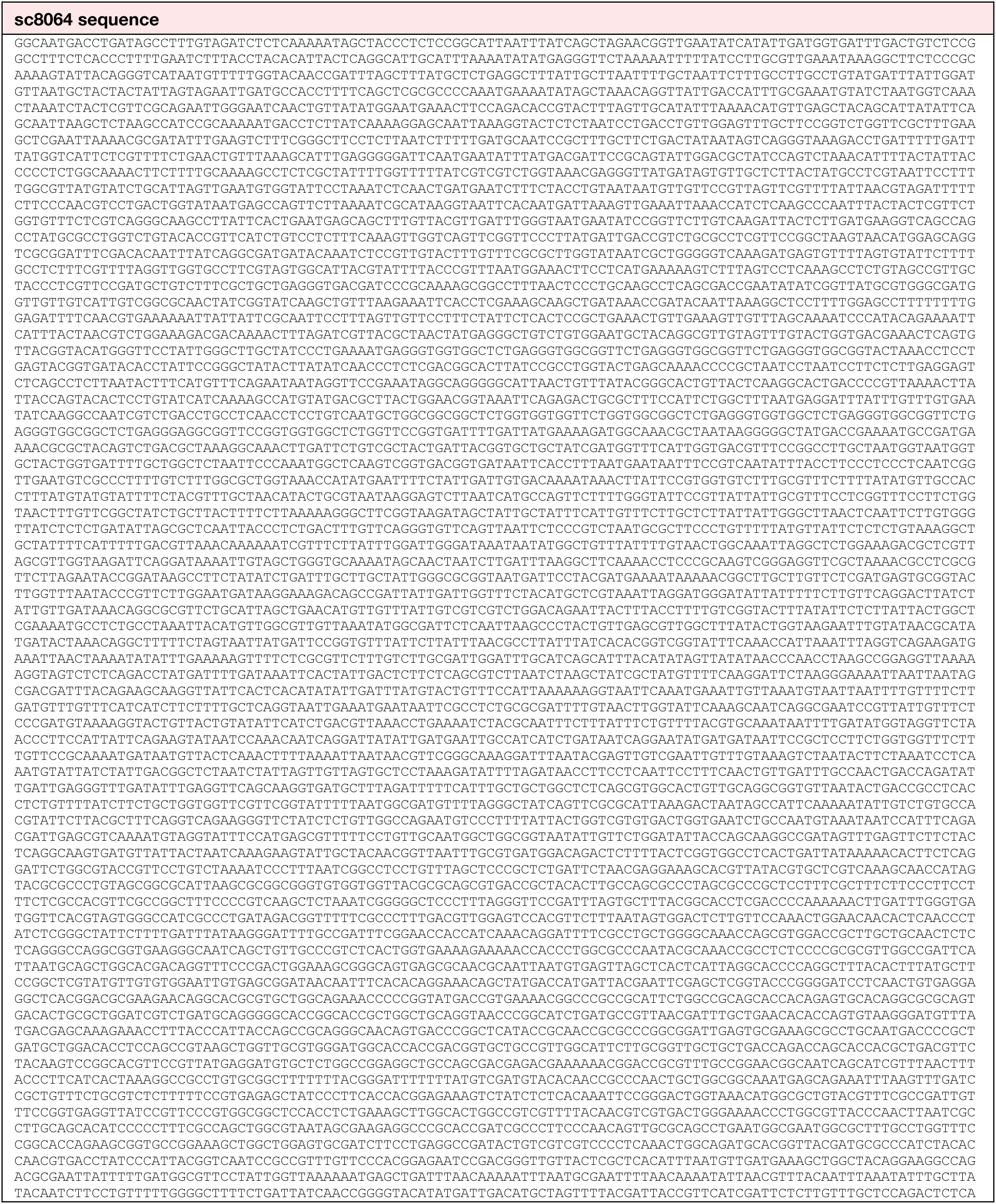
Sequence of scaffold sc8064.

**Supplementary Table 2.** Triangle variants used in this study. For different experiments (see ‘experiment’ and ‘reference figure’ columns), slight variations of the triangular origami structure have been used differing mostly in the sequence, number and positioning of its linker handles. For simplicity, the final staple mixture was divided into three sub-mixtures comprising a constant part (core mix), staples with linker handle extensions on the shell-inner face (internal linker mix), and staples with linker handle extensions on the shell-outer face (external linker mix). For foldings, these mixtures were mixed according to their staple count (equal staple concentrations in the final mixture). The corresponding complementary functionalised oligonucleotides (see Supplementary Table 3 for sequences), if any, for both faces are listed in additional columns. Note that cholesterol-bearing strands were not included in folding reactions but added just before mixing with vesicle solutions (see Materials & Methods section). Refer to the following supplementary tables for sequences. *Linker handles for FLUO1 were left single-stranded (no fluorescence labelling was required).

| experiment | reference figure | core mix | internal linker mix | external linker mix | internal functionalised strands | external functionalised strands |
| --- | --- | --- | --- | --- | --- | --- |
| Vesicle deformation & buds in cryoEM; octahedral DCVs | Fig. 1D<br>S. Fig. 10 | C1 | I1 | E1 | CHOL1 | *no linkers* |
| Budding by tonicity | Fig. 2C &<br>S. Fig. 14 | X6-20 / E6_1<br>C1 | I1 | E1 | CHOL1 | *no linkers* |
| Budding by membrane anchor (b-NAv) | Fig. 2D &<br>S. Fig. 15A | C1 | I3 | E2 | BIOTIN | FLUO1 & bead purification linkers |
| Budding by lipid mixture | Fig. 2E &<br>S. Fig. 16 | X6-20 / E6_1<br>C1 | I1 | E1 | CHOL1 | *no linkers* |
| Budding from phase-separated vesicles | Fig. 2F &<br>S. Fig. 17 | X6_20 /<br>E6_8<br>C1 | I1 | E3 | CHOL1 | FLUO2 |
| Budding kinetics | Fig. 2G &<br>S. Fig. 18 | X6_20/E6_1<br>C1 | I1 | E1 | CHOL1 | *no linkers* |
| Inward budding | Fig. 3B, C | C1 | I4 | E4 | *no linkers* | CHOL1 |
| Bivesicular budding | Fig. 4B | C1 | I1 | E5 | CHOL1 | CHOL2 & FLUO1* |
| Cholesterol position screen | S. Fig. 3 | C1 | I5 or I6 | E1 | CHOL2 or<br>CHOL3 | *no linkers* |
| Cholesterol count screen | S. Fig. 4 | C1 | I7 or I8 or I9 | E6 | CHOL4 | FLUO1 |
| Influence of GVs on triangle assembly | S. Fig. 5 | C1 | I10 or I11 or<br>I12 | E6 | CHOL4 | FLUO1 |
| Influence of GVs on triangle assembly 2 | S. Fig. 6 | X6_2/E6_2<br>C1 | I10 | E6 | CHOL4 | FLUO1 |
| DCVs from triangles with 3/6/9 cholesterol | S. Fig. 8 | X6_2/3/4<br>C1 | I10 or I11 or<br>I12 | E6 | CHOL4 | FLUO1 |
| DNA-lipid colocalisation in agarose gel electrophoresis | S. Fig. 11 | C1 | I10 | E6 | CHOL4 | FLUO1 |
| Assembly as the driving force for DCV budding | S. Fig. 12 | C1 or C2 | I2 | E3 | CHOL2 | FLUO2 |
| Preassembled shells | S. Fig. 13 | C1 | I1 | E1 | CHOL1 | *no linkers* |
| Budding by membrane anchor (chol.) | S. Fig. 15B | C1 | I2 | E2 | CHOL2 | FLUO1 & bead purification linkers |

**Supplementary Table 3.**
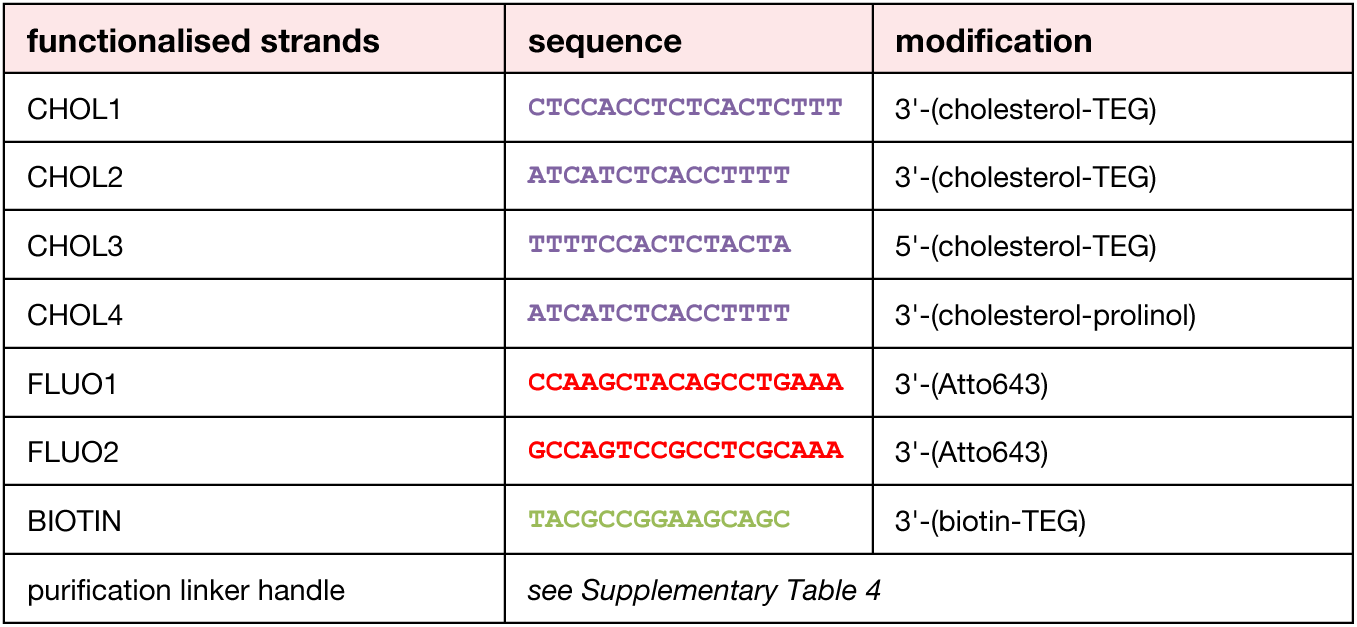
List of functionalised strands.

**Supplementary Table 4.**
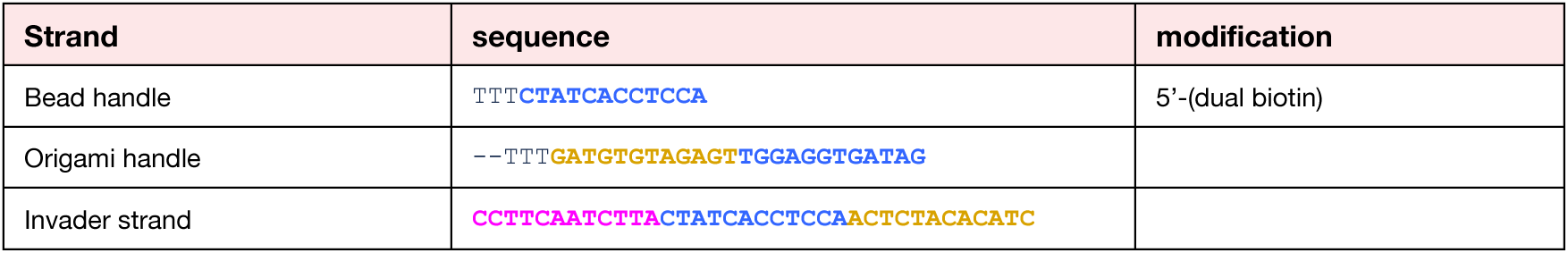
List of sequences for magnetic bead purification of DNA origami. Bead handles are coupled to Dynabeads M-270 Streptavidin. The stated sequence of the origami handle is part of a staple with the full sequences listed in Supplementary Table 20. Complementary sequence snippets are highlighted. The invader strands can optionally be removed from the origami by its reverse complement.

**Supplementary Table 5.**
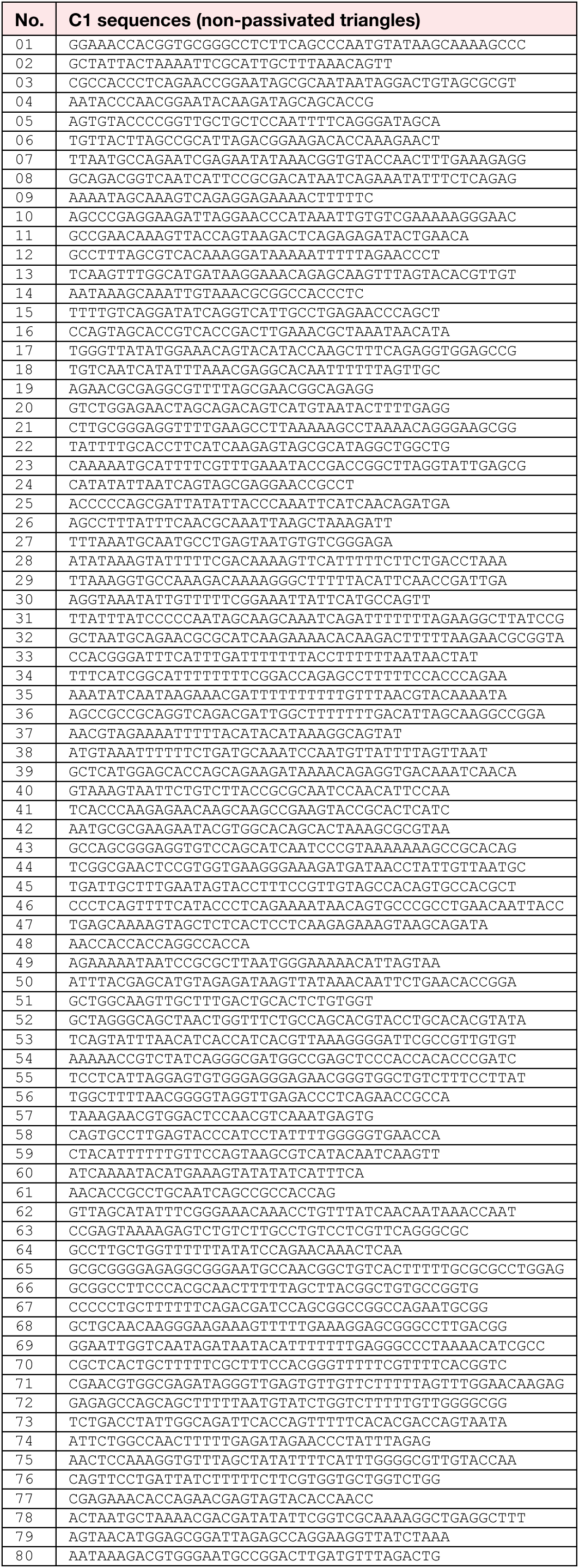
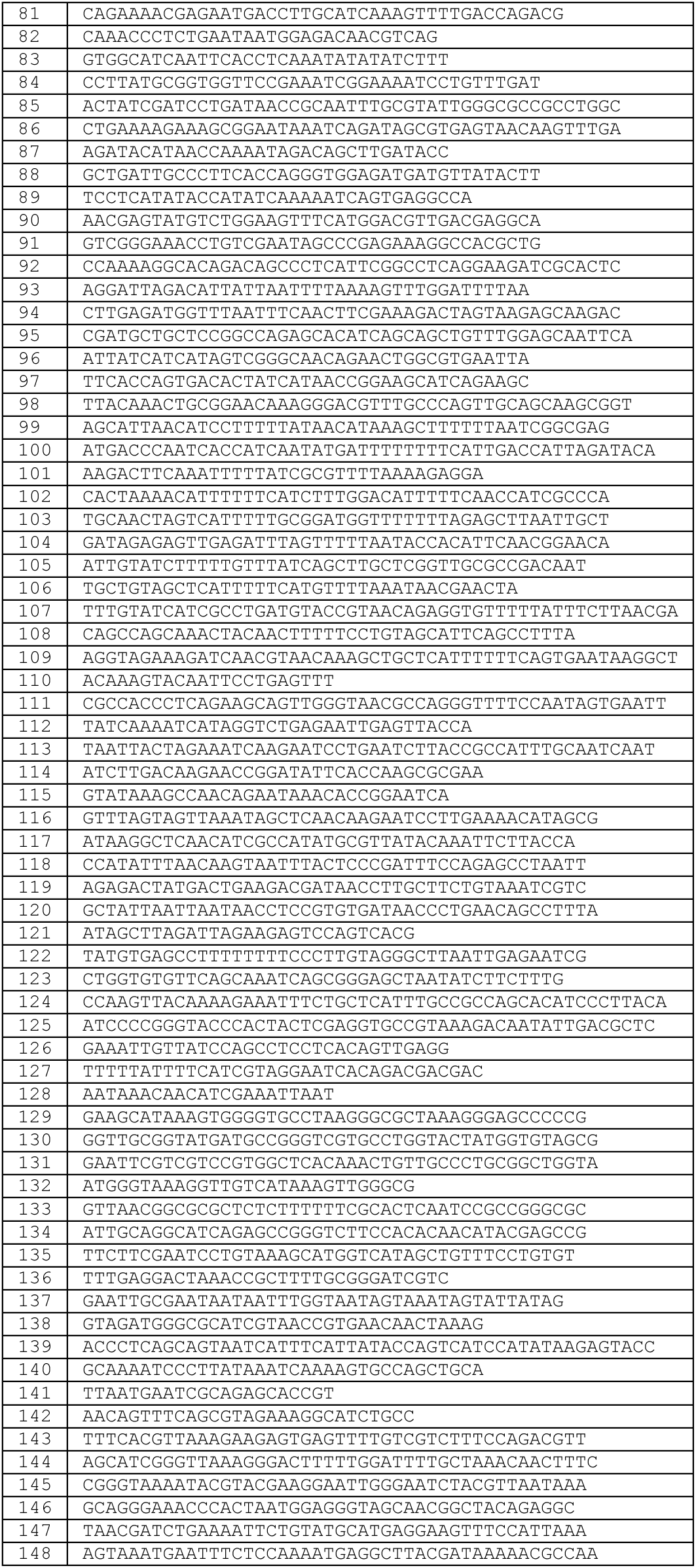
List of staples in core mix 1 (C1). This staple mixture was used to fold non-passivated triangles capable of self-assembly into shells by shape-complementary base-stacking interactions.

**Supplementary Table 6.**
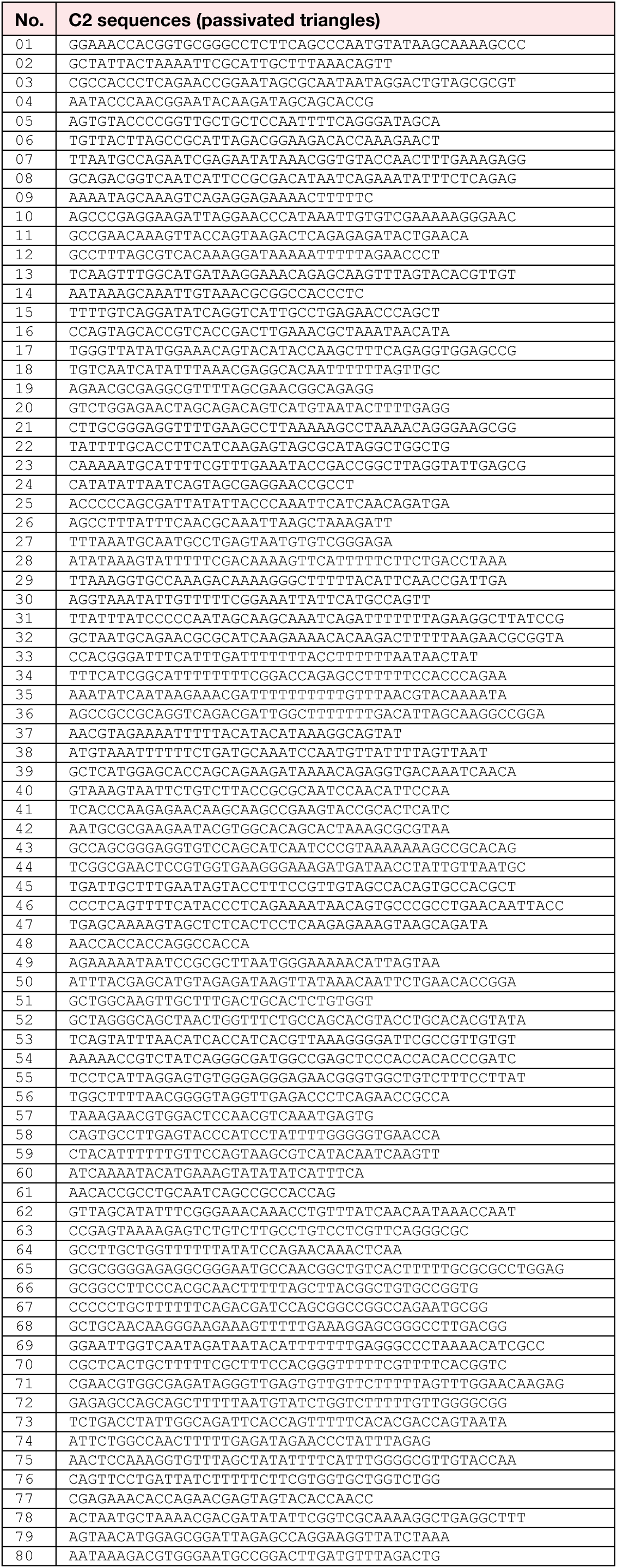
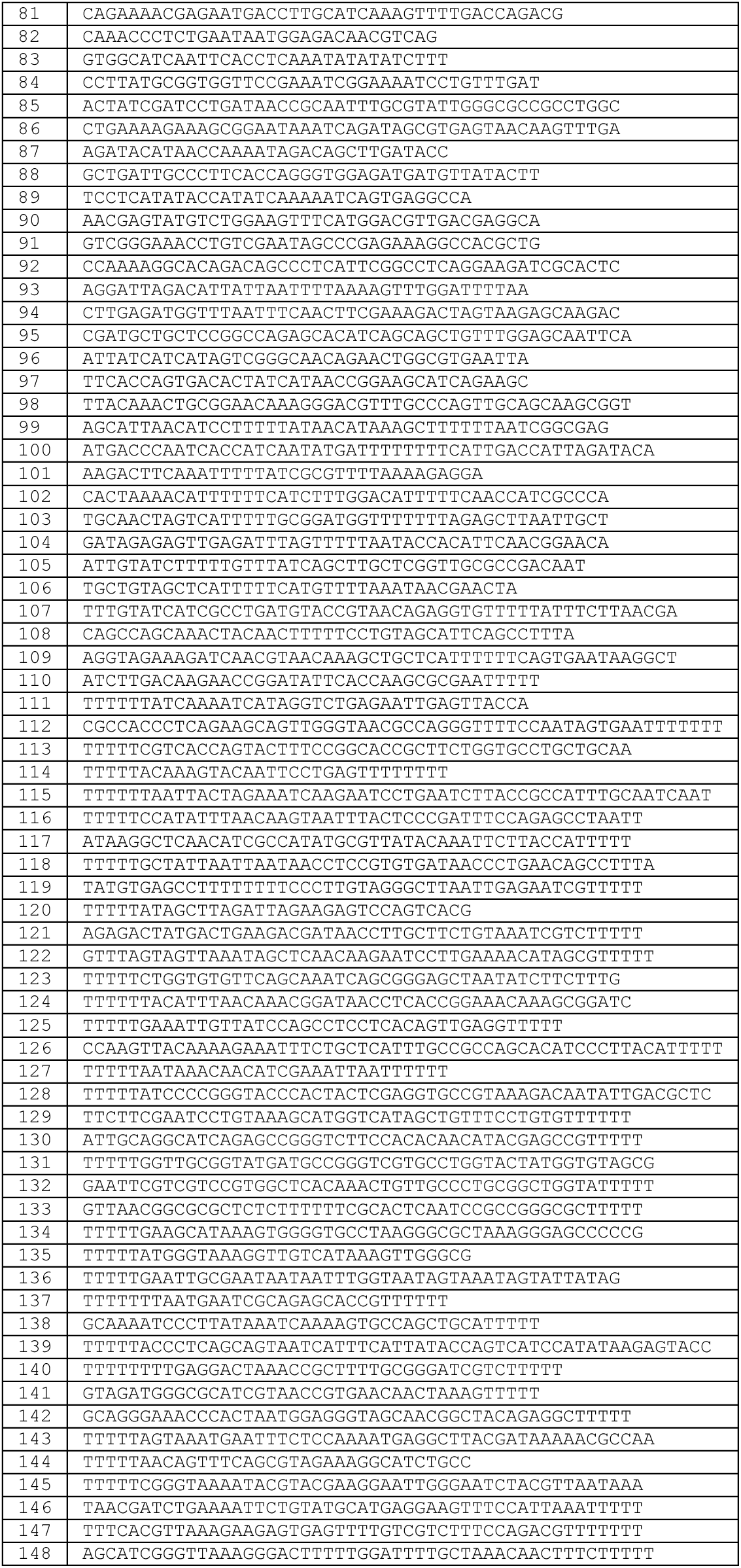
List of staples in core mix 2 (C2). This staple mixture was used to fold passivated triangles incapable of self-assembly into shells by shape-complementary base-stacking interactions due to protruding dT nucleotides at the stacking contacts.

**Supplementary Table 7.**
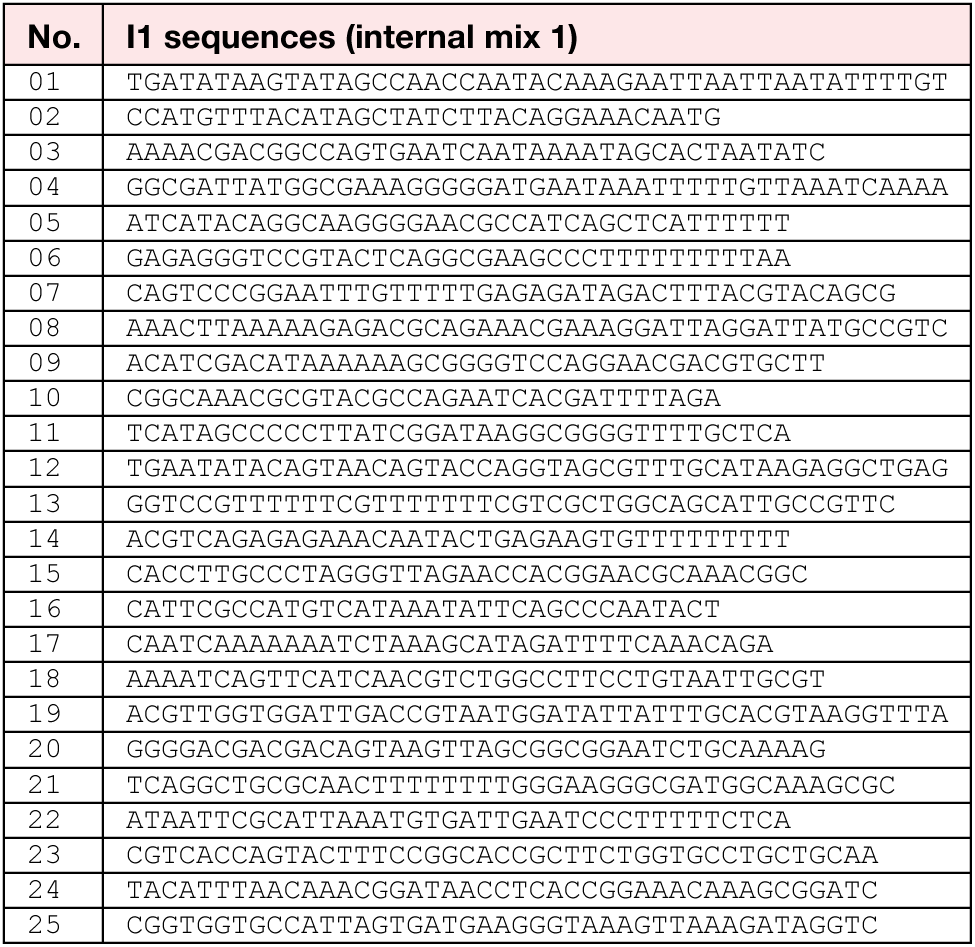
List of staples in internal linker mix 1 (I1).

**Supplementary Table 8.**
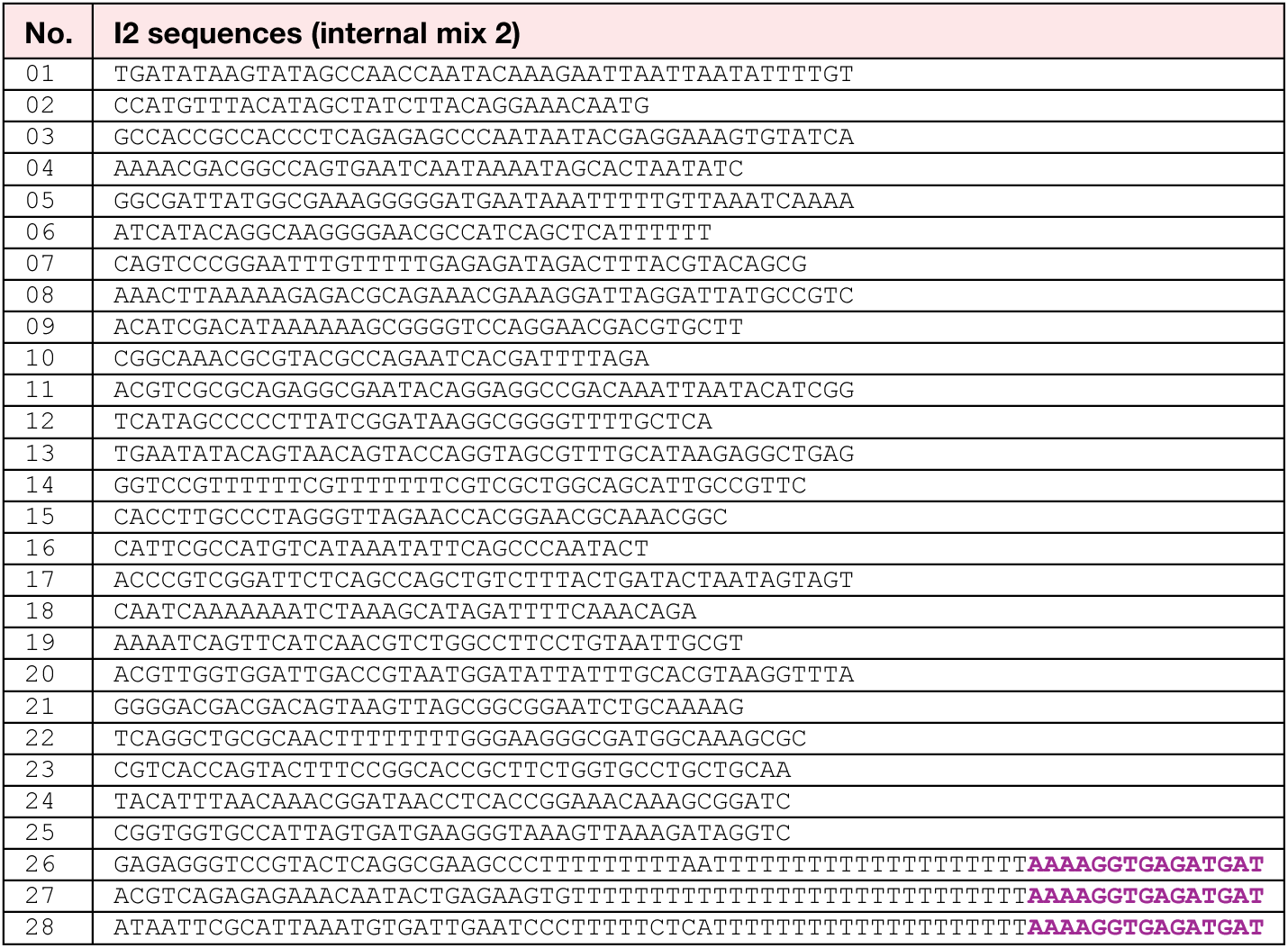
List of staples in internal linker mix 2 (I2). Highlighted sequences are complementary to functionalised oligonucleotides (Supplementary Table 3).

**Supplementary Table 9.**
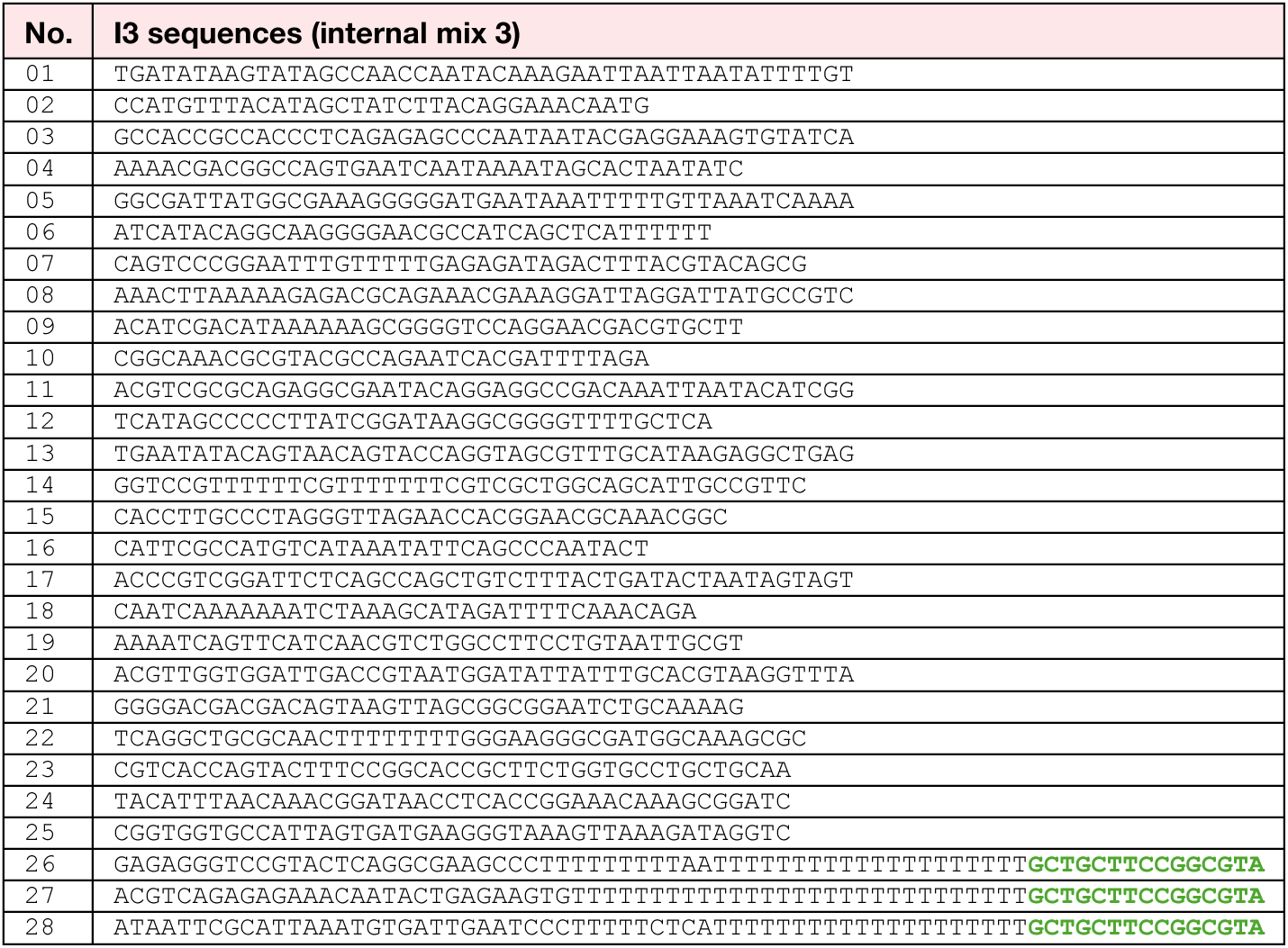
List of staples in internal linker mix 3 (I3). Highlighted sequences are complementary to functionalised oligonucleotides (Supplementary Table 3).

**Supplementary Table 10.**
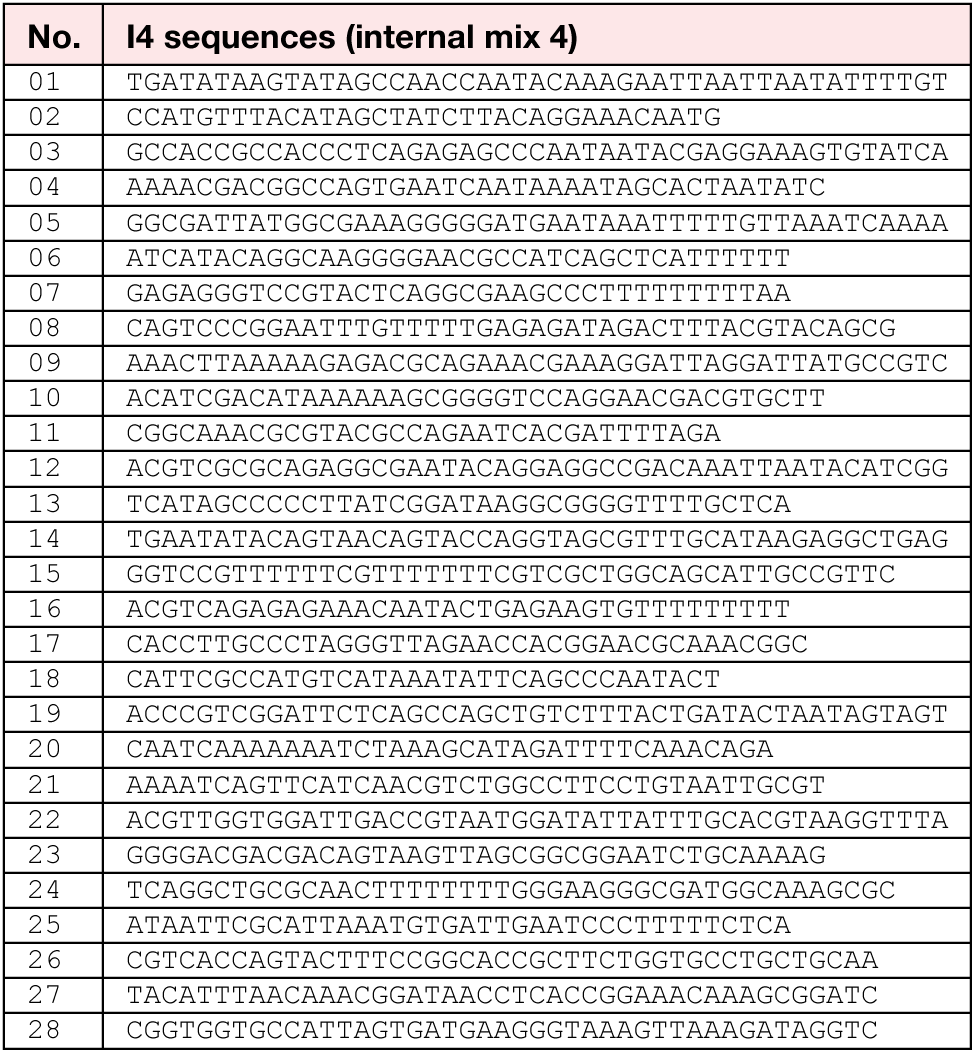
List of staples in internal linker mix 4 (I4).

**Supplementary Table 11.**
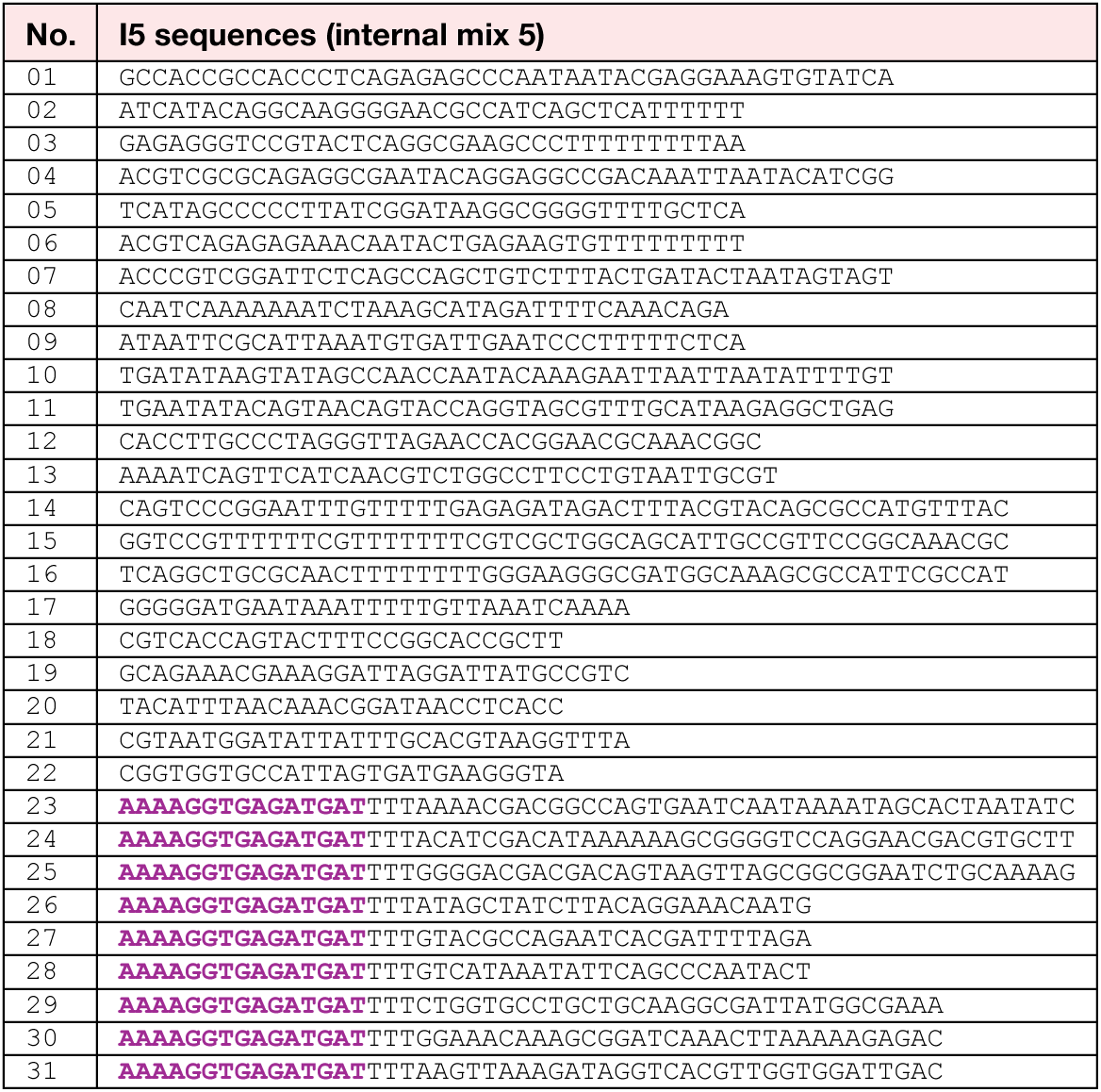
List of staples in internal linker mix 5 (I5). Highlighted sequences are complementary to functionalised oligonucleotides (Supplementary Table 3).

**Supplementary Table 12.**
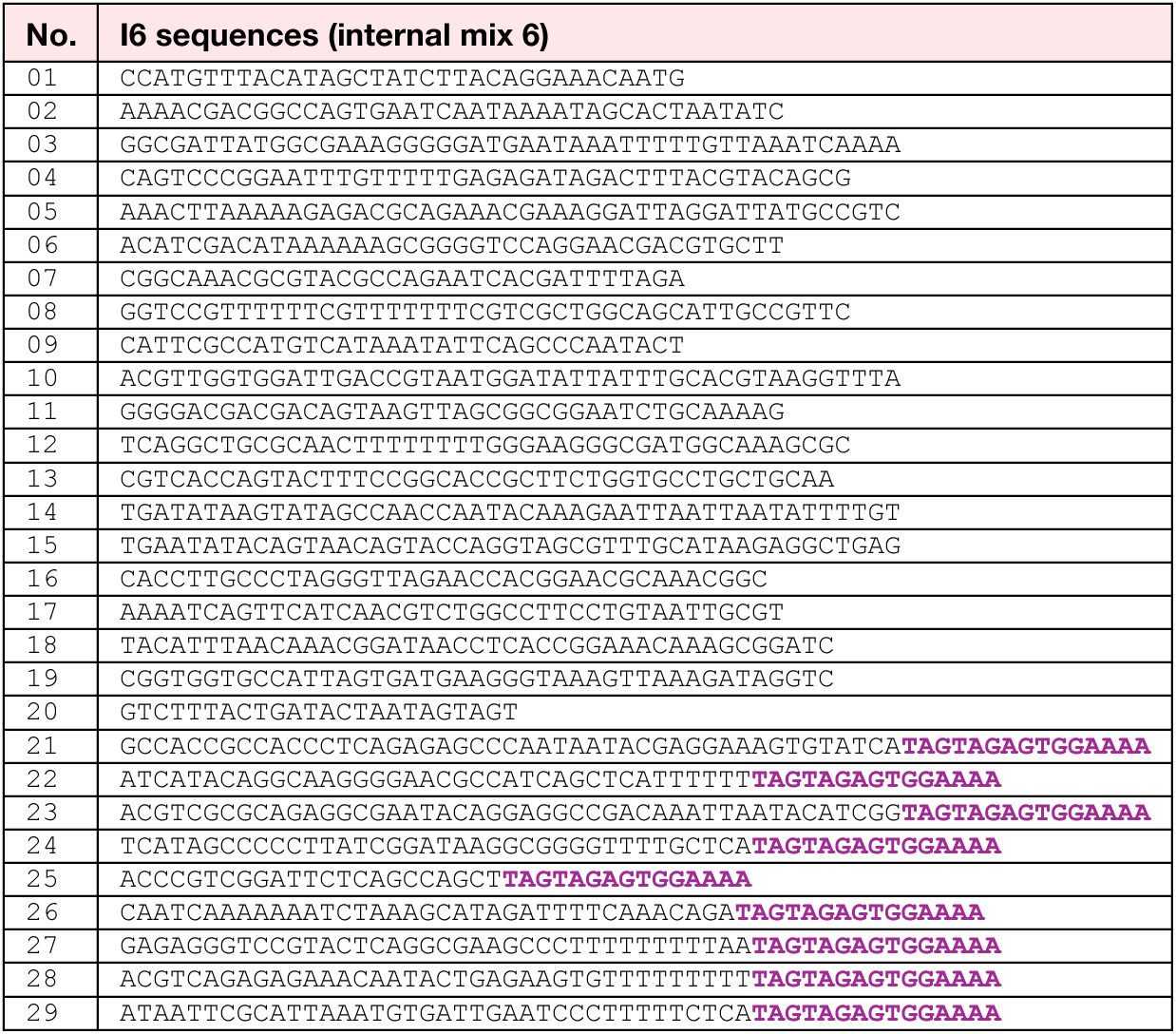
List of staples in internal linker mix 6 (I6). Highlighted sequences are complementary to functionalised oligonucleotides (Supplementary Table 3).

**Supplementary Table 13.**
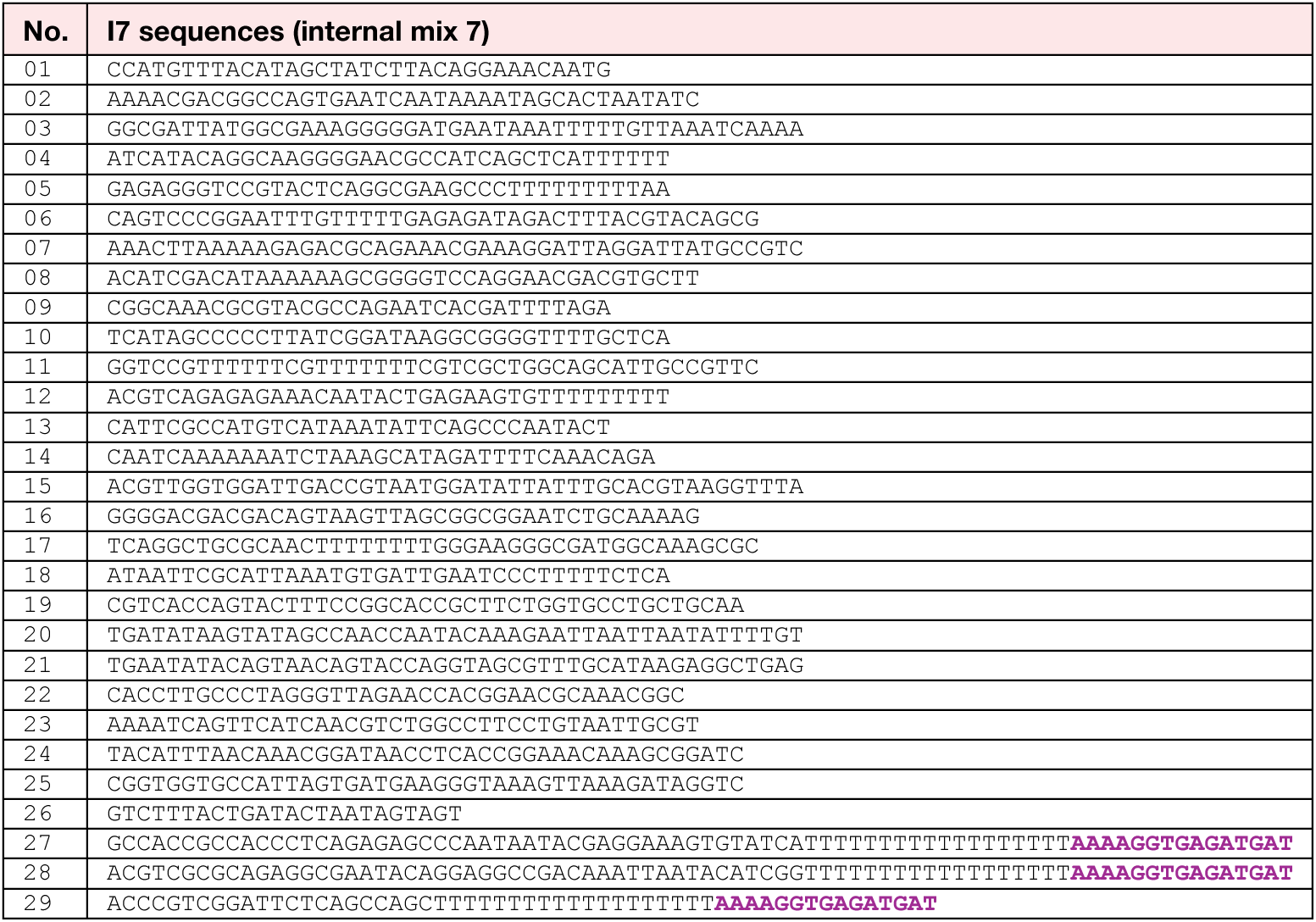
List of staples in internal linker mix 7 (I7). Highlighted sequences are complementary to functionalised oligonucleotides (Supplementary Table 3).

**Supplementary Table 14.**
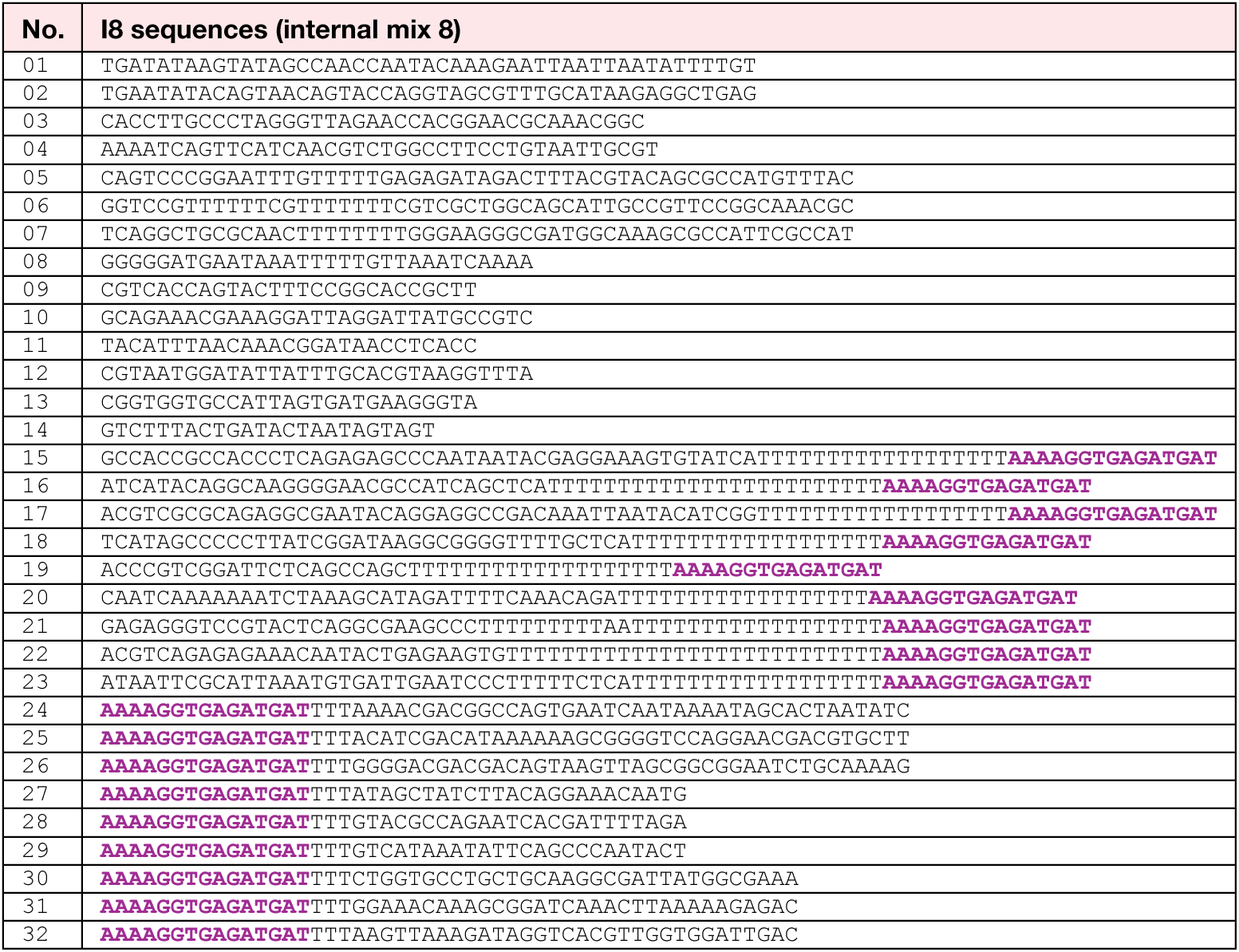
List of staples in internal linker mix 8 (I8). Highlighted sequences are complementary to functionalised oligonucleotides (Supplementary Table 3).

**Supplementary Table 15.**
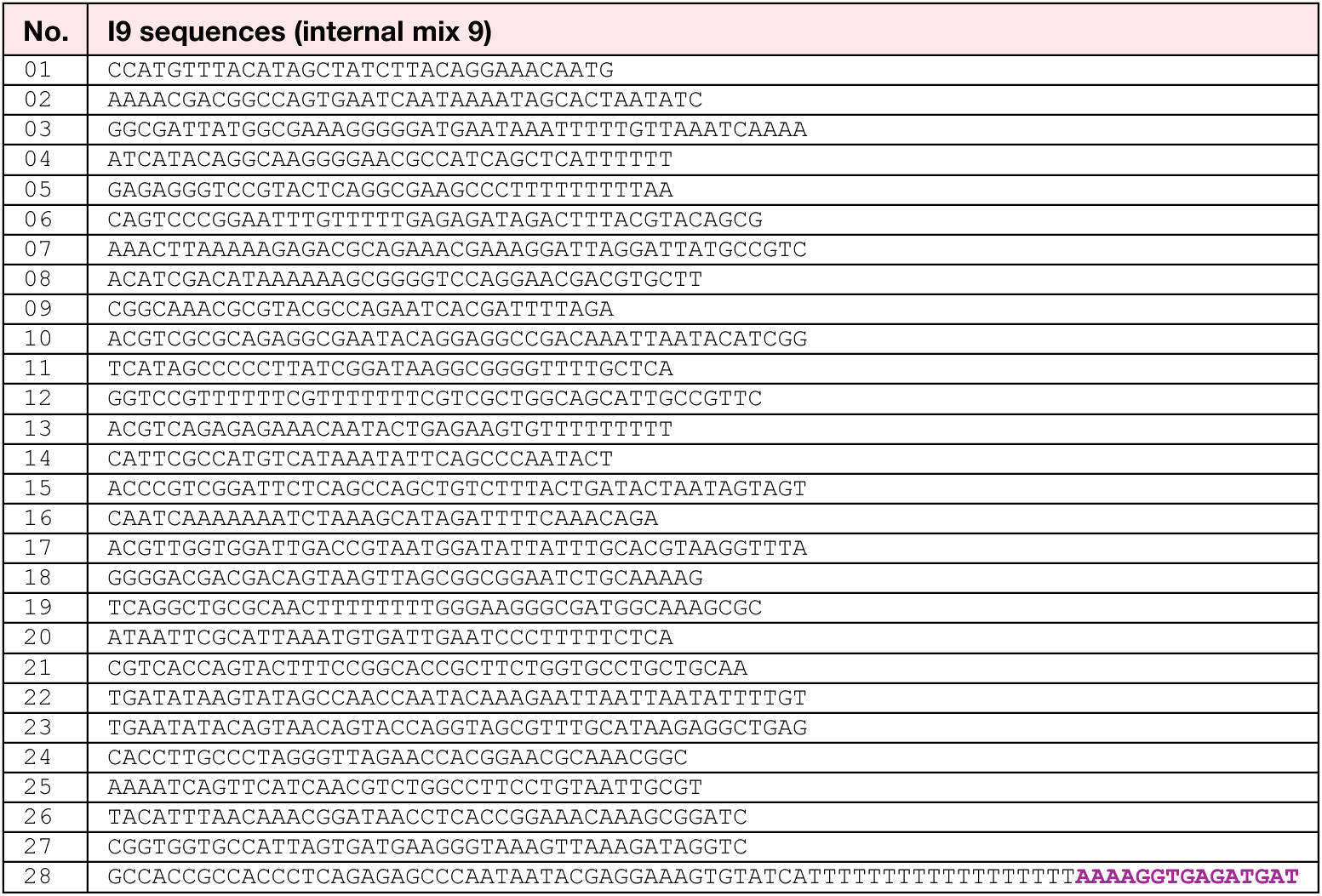
List of staples in internal linker mix 9 (I9). Highlighted sequences are complementary to functionalised oligonucleotides (Supplementary Table 3).

**Supplementary Table 16.**
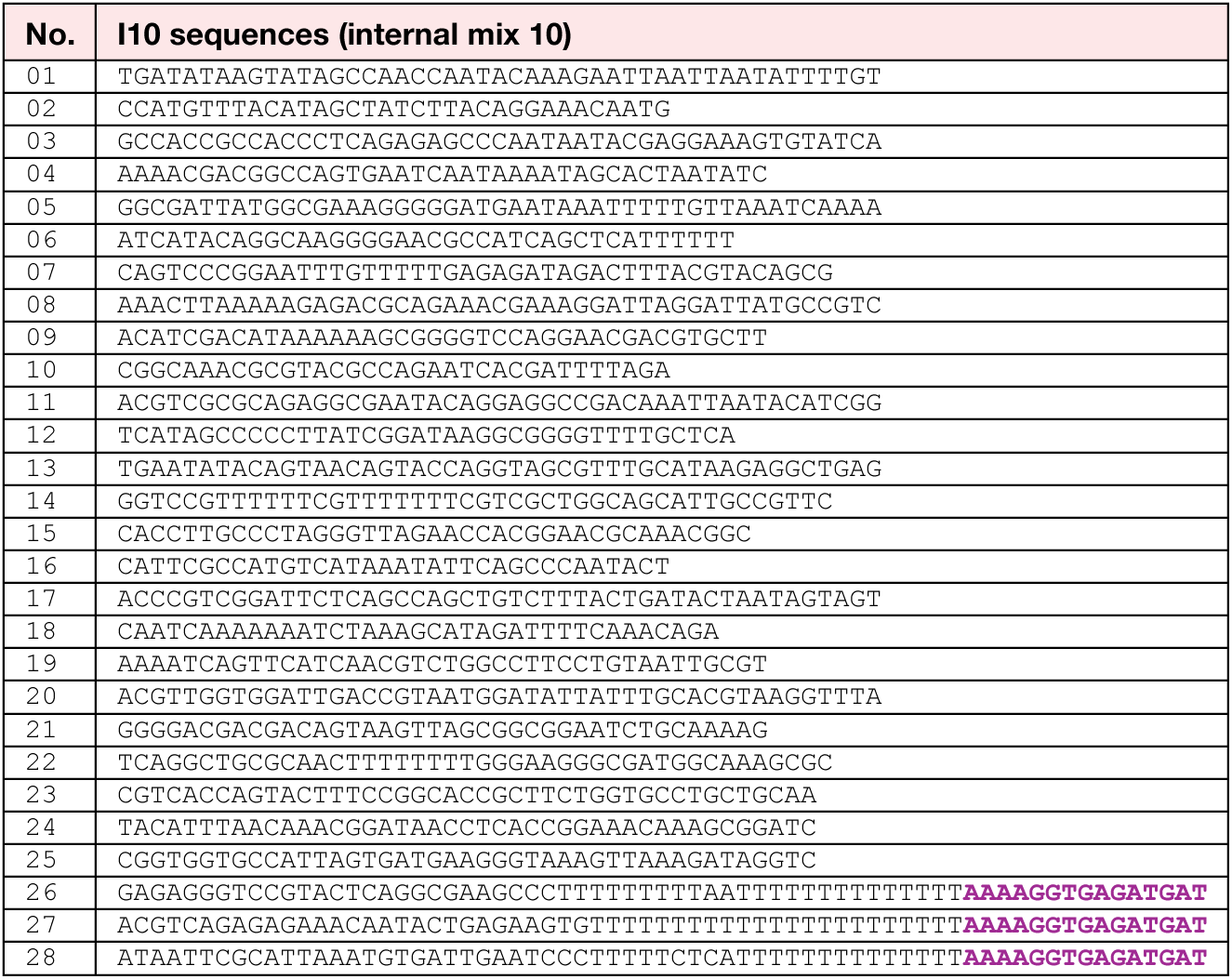
List of staples in internal linker mix 10 (I10). Highlighted sequences are complementary to functionalised oligonucleotides (Supplementary Table 3).

**Supplementary Table 17.**
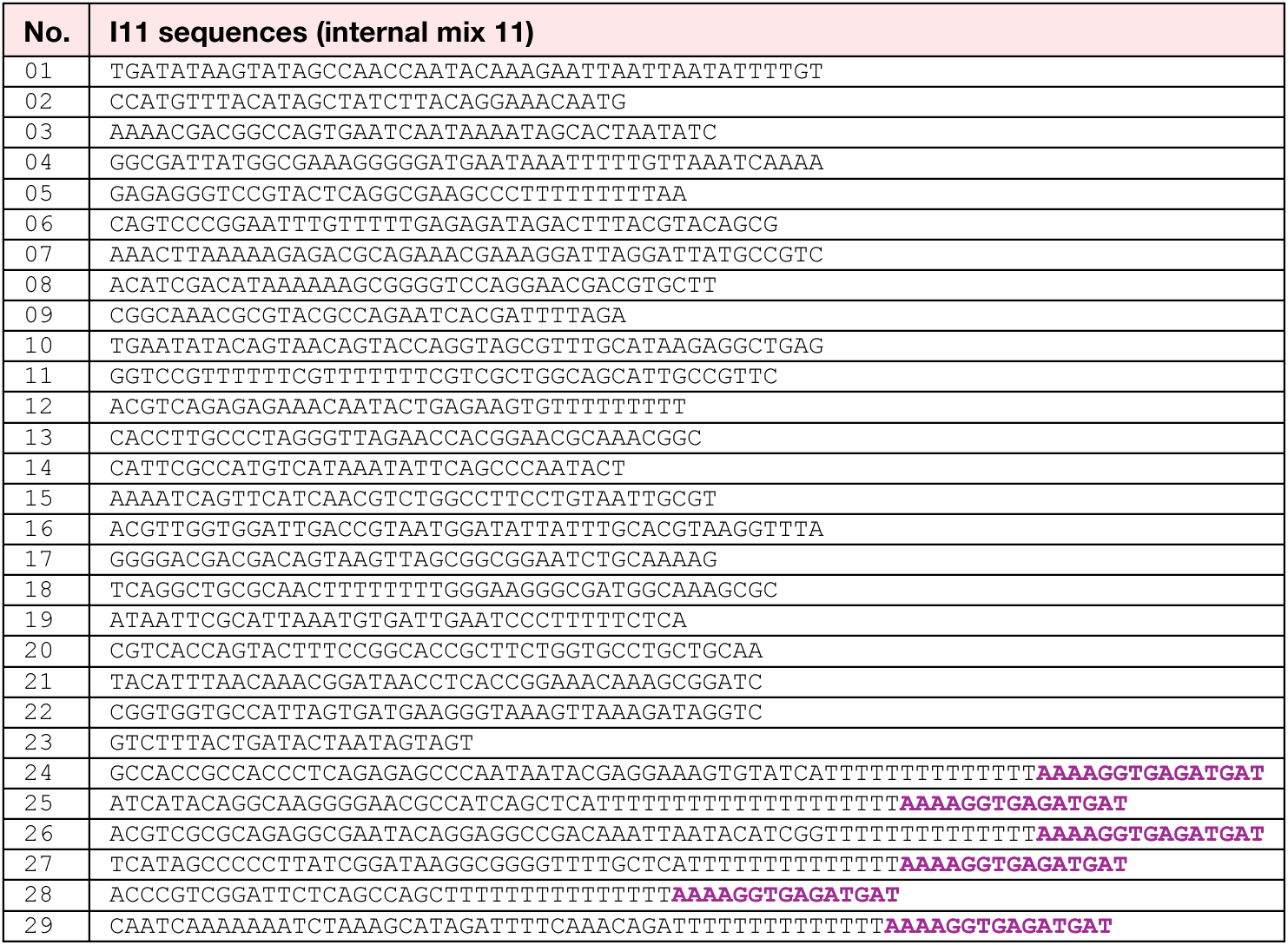
List of staples in internal linker mix 11 (I11). Highlighted sequences are complementary to functionalised oligonucleotides (Supplementary Table 3).

**Supplementary Table 18.**
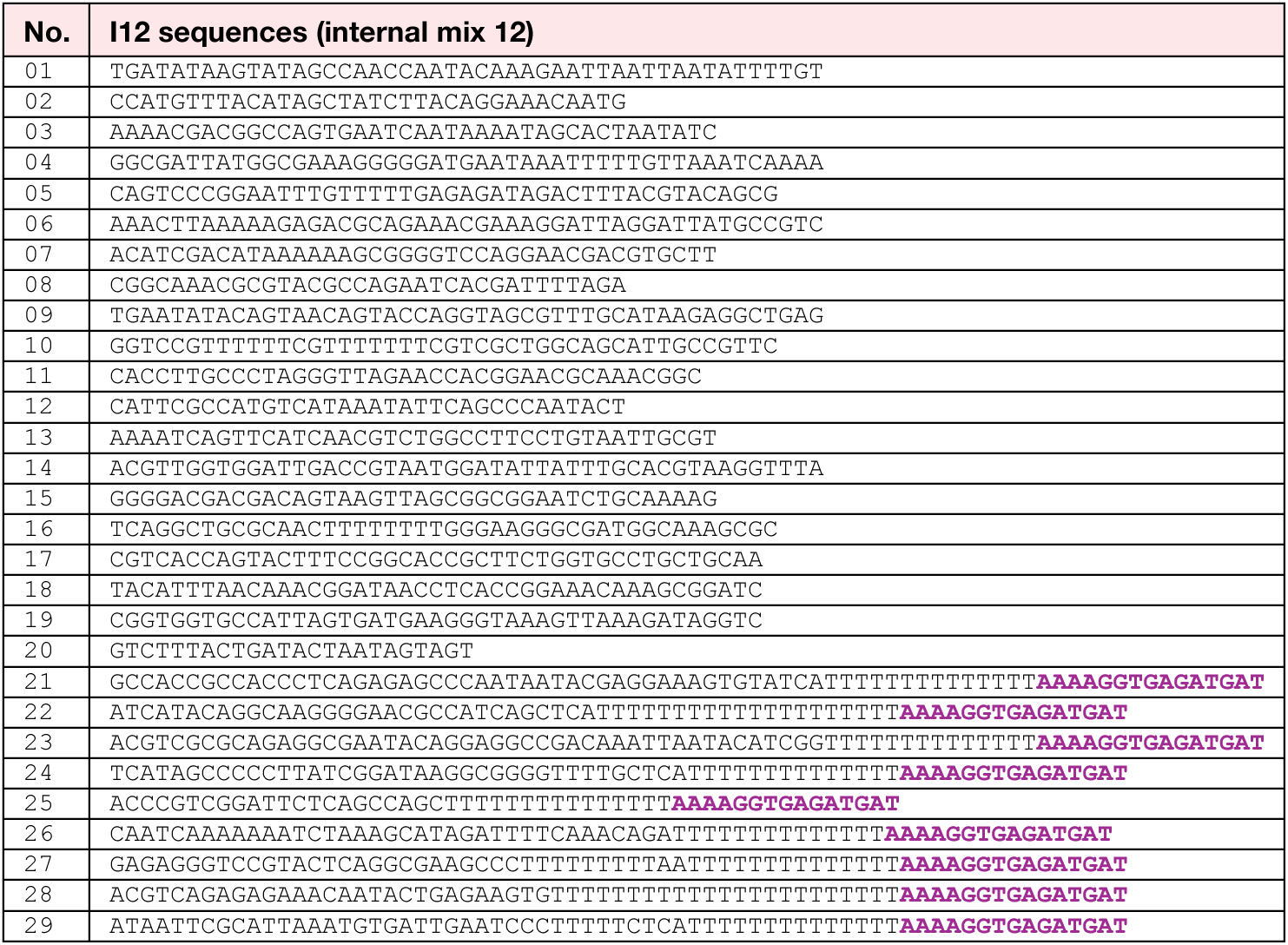
List of staples in internal linker mix 12 (I12). Highlighted sequences are complementary to functionalised oligonucleotides (Supplementary Table 3).

**Supplementary Table 19.**
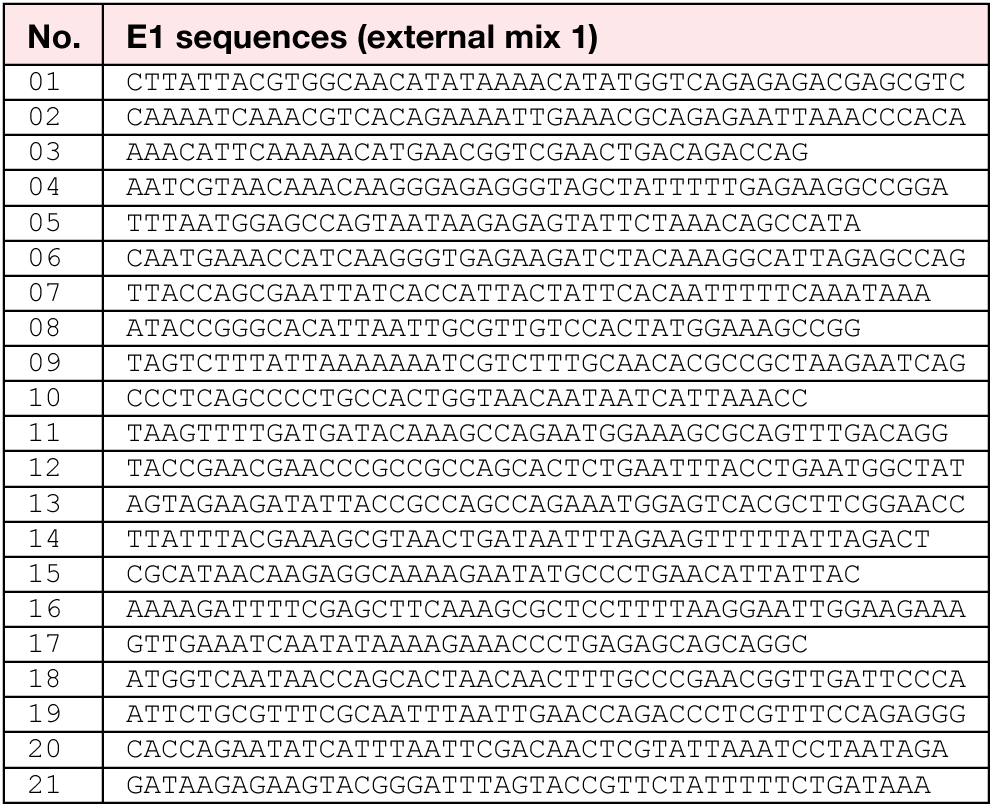
List of staples in external linker mix 1 (E1).

**Supplementary Table 20.**
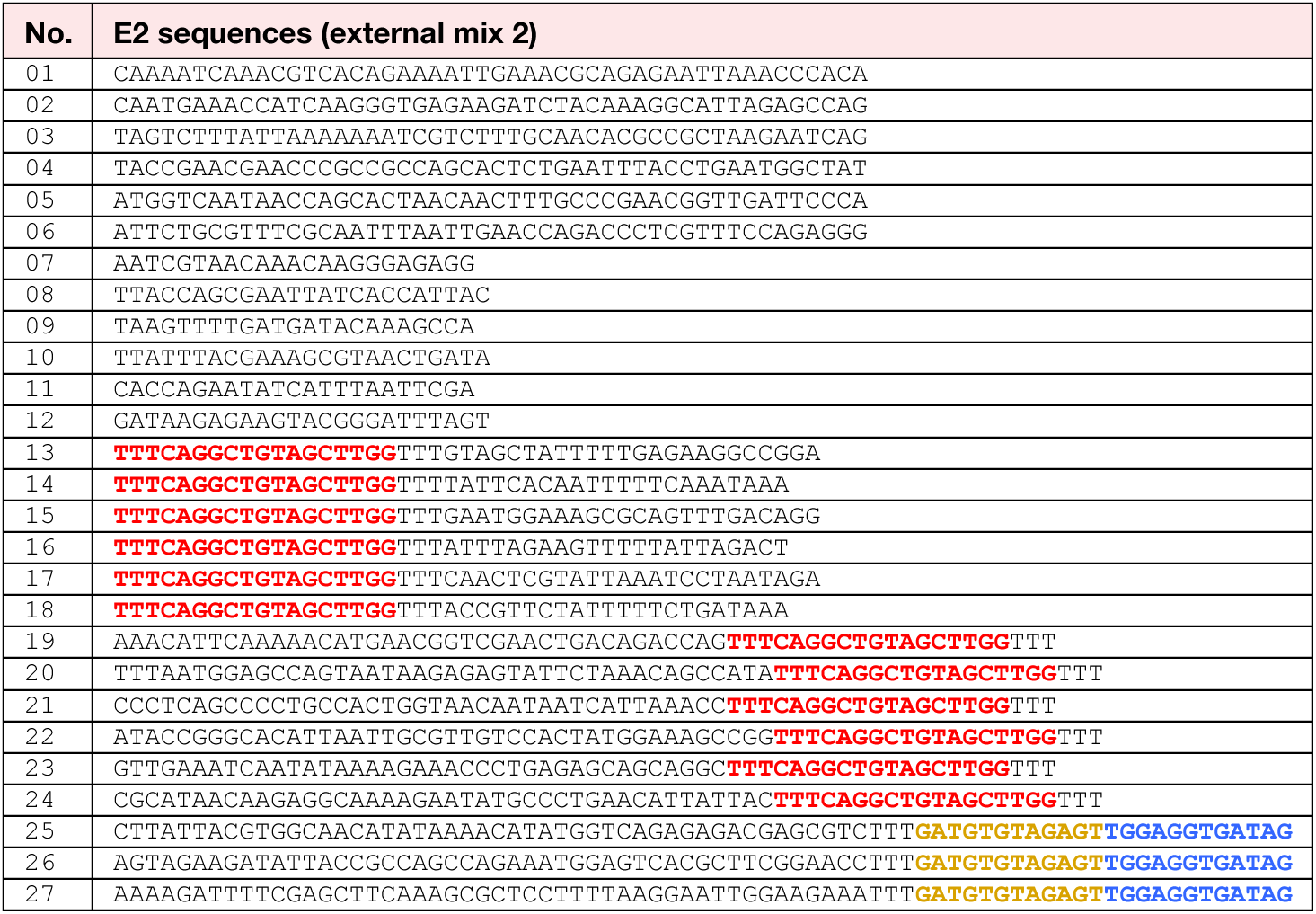
List of staples in external linker mix 2 (E2). Highlighted sequences are complementary to functionalised oligonucleotides (Supplementary Tables 3 & 4).

**Supplementary Table 21.**
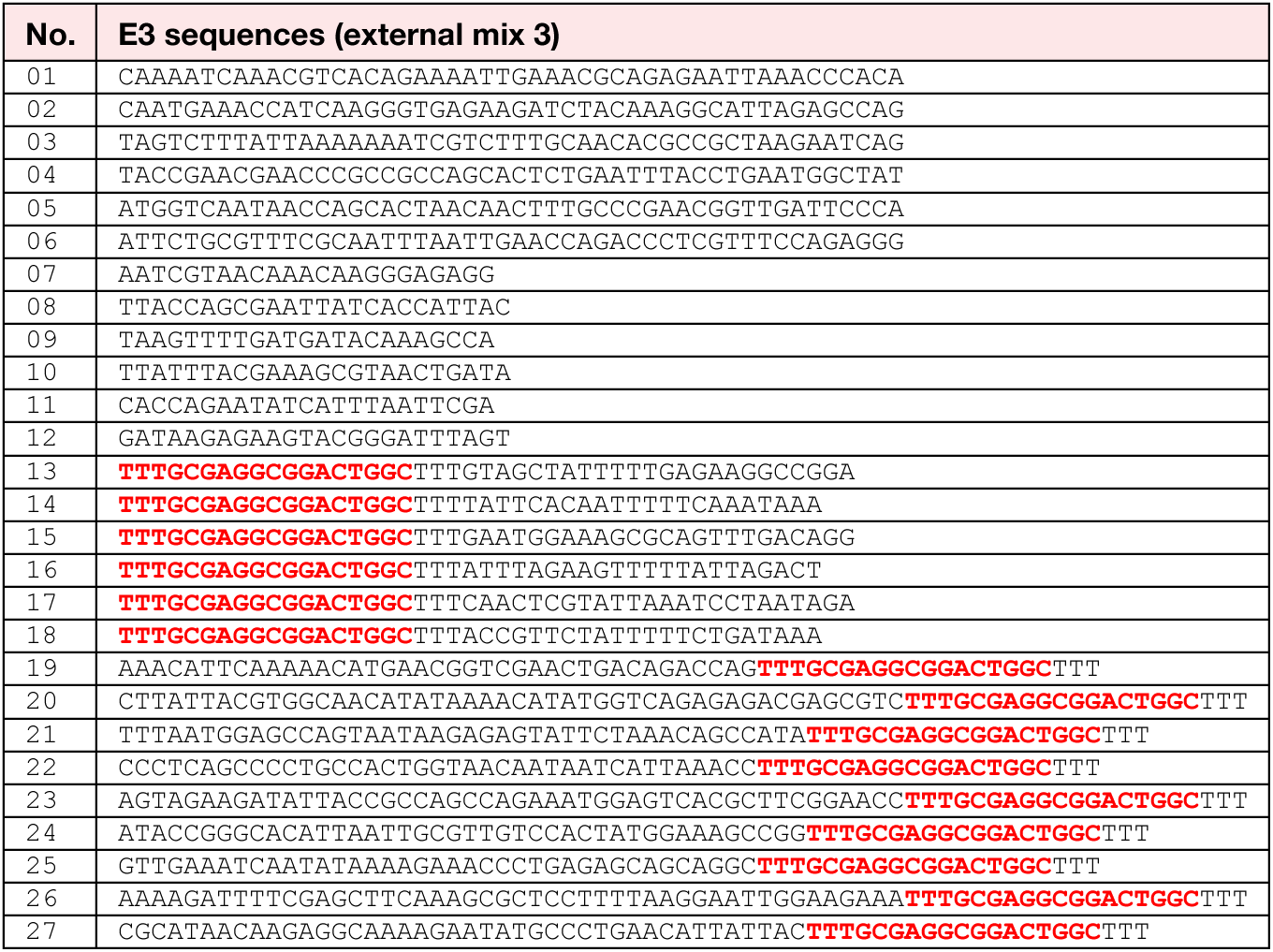
List of staples in external linker mix 3 (E3). Highlighted sequences are complementary to functionalised oligonucleotides (Supplementary Table 3).

**Supplementary Table 22.**
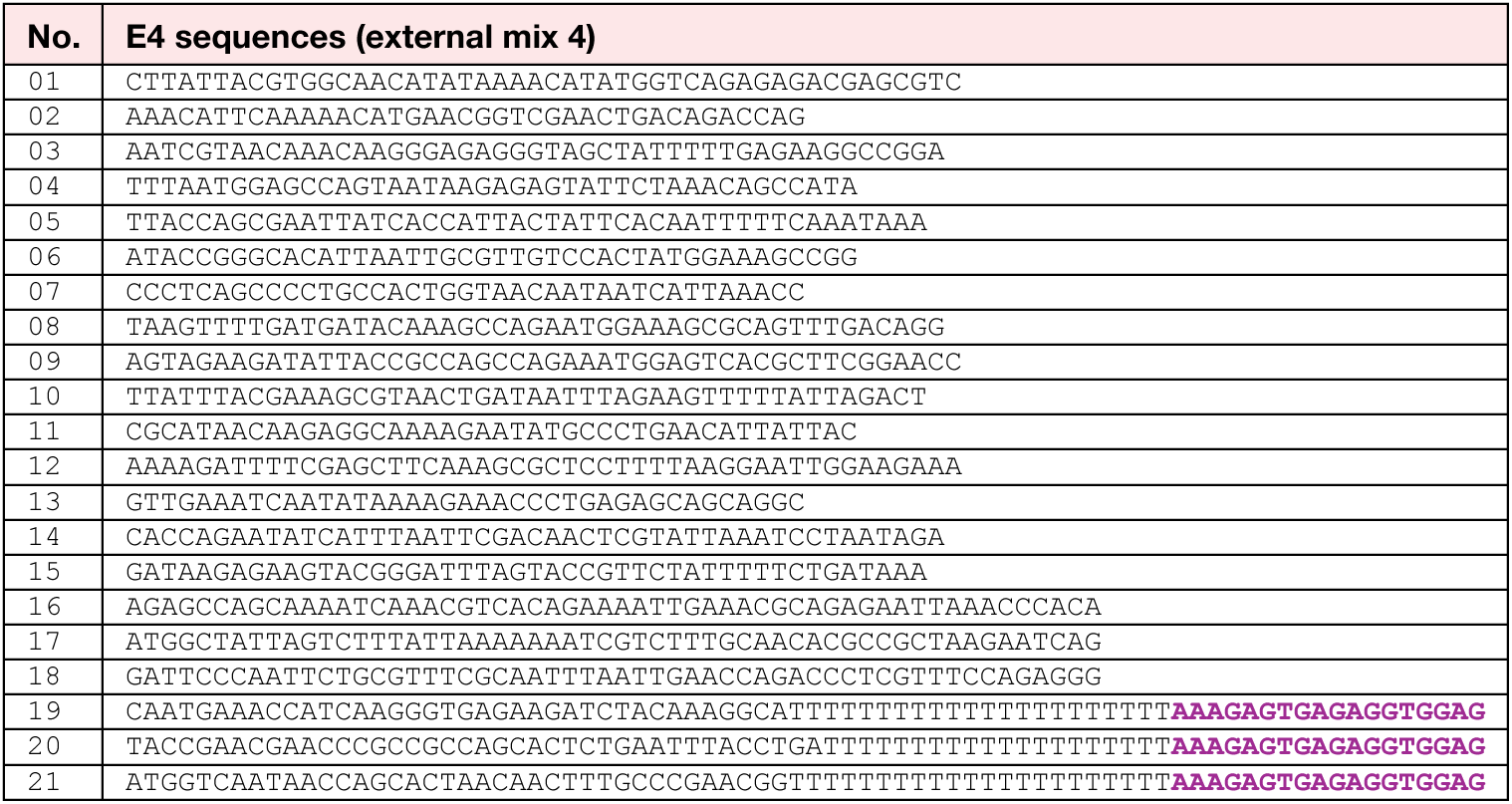
List of staples in external linker mix 4 (E4). Highlighted sequences are complementary to functionalised oligonucleotides (Supplementary Table 3).

**Supplementary Table 23.**
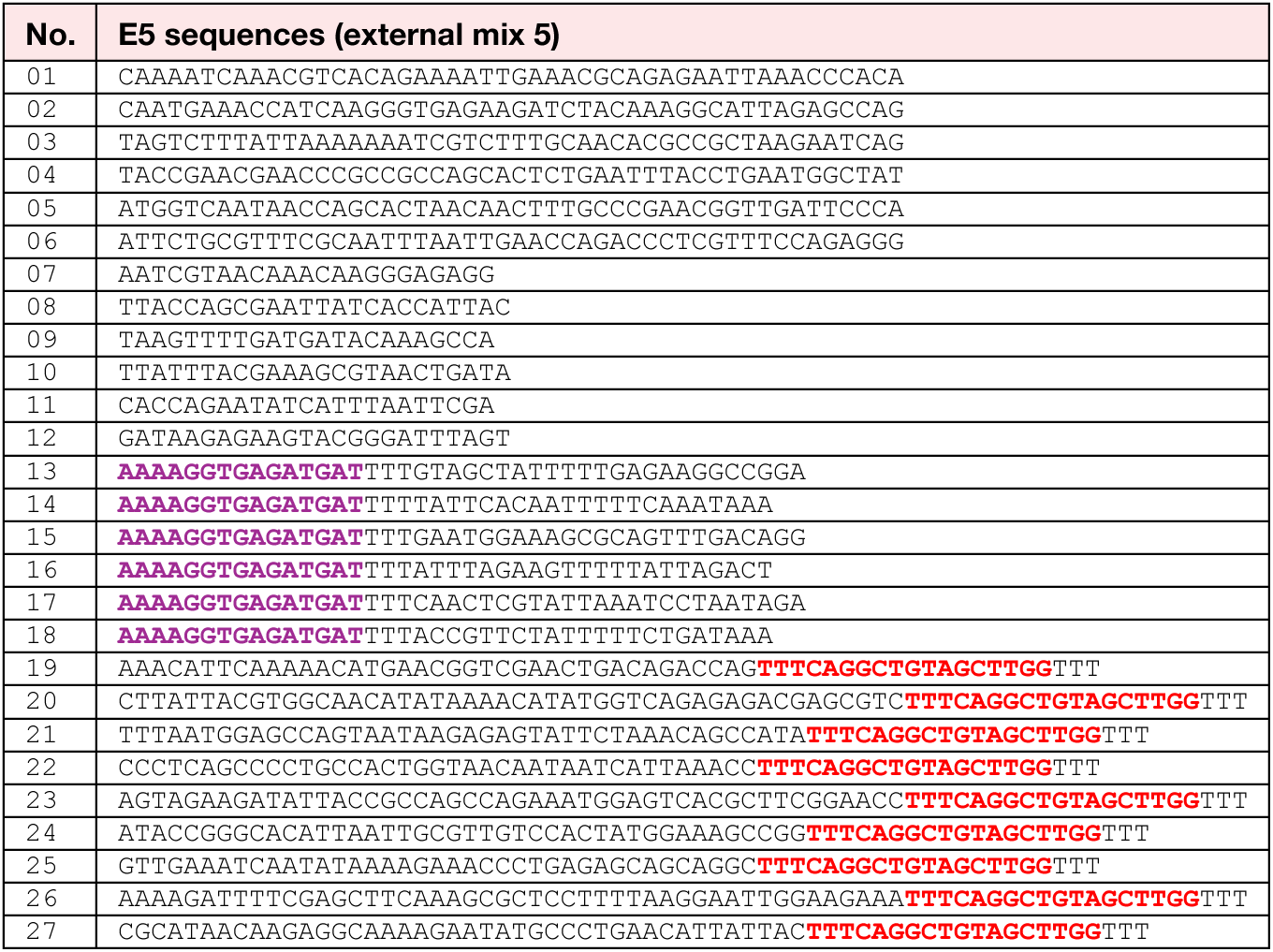
List of staples in external linker mix 5 (E5). Highlighted sequences are complementary to functionalised oligonucleotides (Supplementary Table 3).

**Supplementary Table 24.**
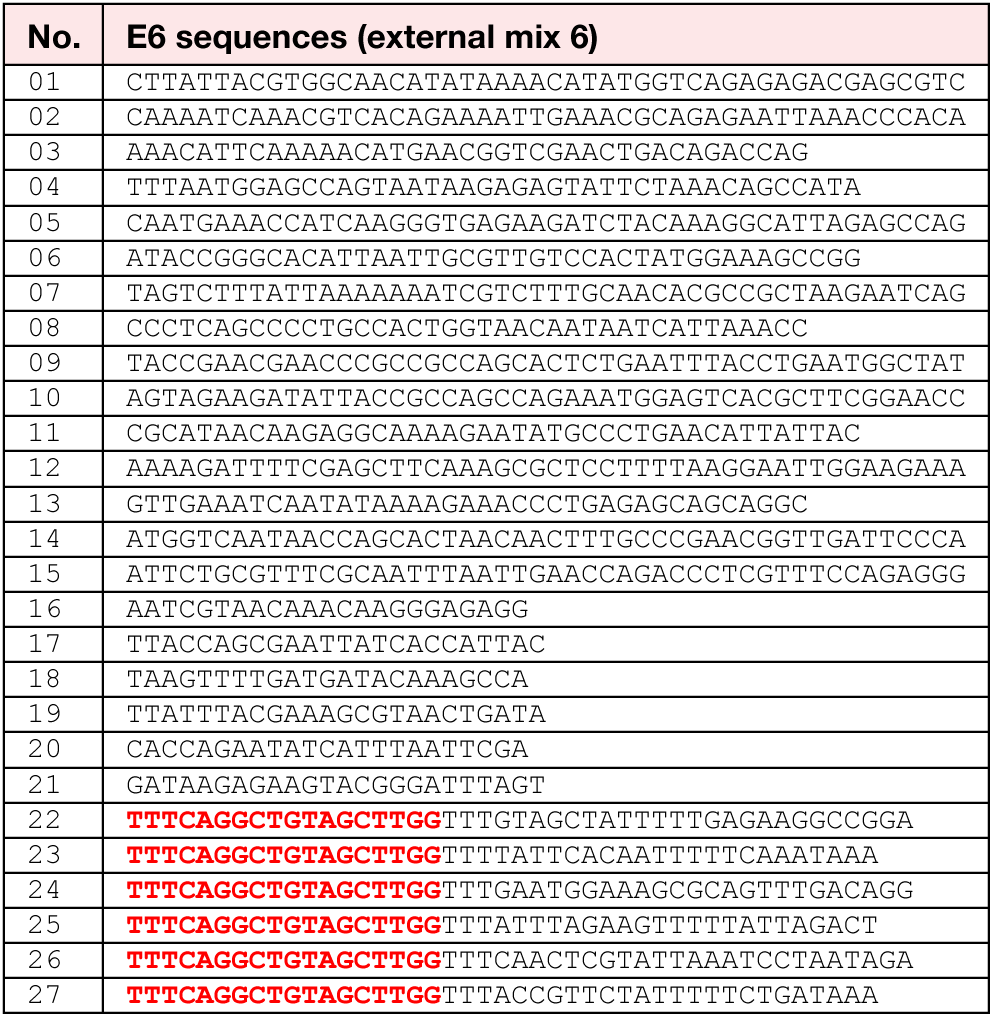
List of staples in external linker mix 6 (E6). Highlighted sequences are complementary to functionalised oligonucleotides (Supplementary Table 3).

**Supplementary Table 25.** Tonicity screen sample compositions. Composition of the samples shown in Fig. 2c. The magnesium optimum for triangle assembly increased with increasing glycine concentrations. Concentrations above or below the optima presented in this table result in incomplete or impaired assembly.

| Osmolality difference | Glycine (mM) | MgCl<sub>2</sub> (mM) | NaCl (mM) | Total volume (μl) |
| --- | --- | --- | --- | --- |
| -78% | 0 | 25 | 26.3 | 30 |
| -58% | 100 | 30 | 26.3 | 30 |
| -42% | 200 | 35 | 26.3 | 30 |
| -22% | 300 | 45 | 26.3 | 30 |
| -4 | 400 | 50 | 26.3 | 30 |
| +12% | 500 | 55 | 26.3 | 30 |
| +33% | 600 | 65 | 26.3 | 30 |
| +54% | 700 | 75 | 26.3 | 30 |

**Supplementary Table 26.** Buffer compositions.

|  | TRIS<br>(mM) | EDTA<br>(mM) | MgCl<sub>2</sub><br>(mM) | NaCl<br>(mM) | CsCl<br>(mM) | Sucrose<br>(mM) | Glucose<br>(mM) | Glycine<br>(mM) | pH |
| --- | --- | --- | --- | --- | --- | --- | --- | --- | --- |
| Folding Buffer | 5 | 1 | 20 | 5 |  |  |  |  | 8.0 |
| Sodium Buffer | 5 | 1 | 5 | 305 |  |  |  |  | 8.0 |
| Caesium Buffer | 5 | 1 | 5 | 5 | 320 |  |  |  | 8.0 |
| Sucrose Buffer | 10 | 1 |  |  |  | 470 |  |  | 8.0 |
| Imaging Buffer A | 5 | 1 | 5 | 5 |  | 280 |  |  | 8.0 |
| Imaging Buffer B | 5 | 1 | 5 | 5 |  |  | 280 |  | 8.0 |
| Isotonic assembly buffer | 5 | 1 | 210 | 5 |  |  |  |  | 8.0 |
| Glycine buffer | 5 | 1 | 5 |  |  |  |  | 1000 | 6.8 |

